# Evolutionary dynamics of the SKN-1 → MED → END-1,3 regulatory gene cascade in *Caenorhabditis* endoderm specification

**DOI:** 10.1101/769760

**Authors:** Morris F. Maduro

## Abstract

Gene regulatory networks (GRNs) with GATA factors are important in animal development, and evolution of such networks is an important problem in the field. In the nematode, *Caenorhabditis elegans*, the endoderm (gut) is generated from a single embryonic precursor, E. The gut is specified by an essential cascade of transcription factors in a GRN, with the maternal factor SKN-1 at the top, activating expression of the redundant *med-1,2* divergent GATA factor genes, with the combination of all three contributing to activation of the paralogous *end-3* and *end-1* canonical GATA factor genes. In turn, these factors activate the GATA factors genes *elt-2* and *elt-7* to regulate intestinal fate. In this work, genome sequences from over two dozen species within the *Caenorhabditis* genus are used to identify putative orthologous genes encoding the MED and END-1,3 factors. The predictions are validated by comparison of gene structure, protein conservation, and putative *cis-*regulatory sites. The results show that all three factors occur together, but only within the Elegans supergroup of related species. While all three factors share similar DNA-binding domains, the MED factors are the most diverse as a group and exhibit unexpectedly high gene amplifications, while the END-1 orthologs are highly conserved and share additional extended regions of conservation not found in the other GATA factors. The MEME algorithm identified both known and previously unrecognized *cis-*regulatory motifs. The results suggest that all three genes originated at the base of the Elegans supergroup and became fixed as an essential embryonic gene regulatory network with several conserved features, although each of the three factors is under different evolutionary constraints. Based on the results, a model for the origin and evolution of the network is proposed. The set of identified MED, END-3 and END-1 factors form a robust set of factors defining an essential embryonic gene network that has been conserved for tens of millions of years, that will serve as a basis for future studies of GRN evolution.

## INTRODUCTION

Central to the development of a metazoan is the activation of tissue-specific gene regulatory networks (GRNs) that drive subdivision of progenitors and emergence of features of terminal differentiation (Davidson 2010). On evolutionary time scales, changes in such networks drive appearance of novel features, but these changes can also occur without changes in morphology or development (Peter and Davidson 2016). Such differences in GRNs that nonetheless drive homologous developmental processes exemplify Developmental System Drift (DSD) (True and Haag 2001). In the nematode genus *Caenorhabditis*, which includes the well-studied species *C. elegans*, examples of DSD include the gene networks that produce the derived character of hermaphroditism, which evolved at least three independent times in the genus, and vulval development (Ellis and Lin 2014; Felix 2007; Haag *et al.* 2018).

A relatively understudied area in *Caenorhabditis* is the evolutionary dynamics of GRNs that drive embryonic development. One reason may be that the close relatives to *C. elegans* exhibit indistinguishable embryogenesis, differing perhaps by the timing of some developmental milestones (Levin *et al.* 2012; Memar *et al.* 2019; Zhao *et al.* 2008). Another reason for the paucity of evo-devo studies in embryogenesis is that the dissection of a GRN requires cause-and-effect associations to be probed through experimental perturbations (Davidson *et al.* 2002). The powerful tools of forward and reverse genetics in *C. elegans* have only recently become available in related species, most notably *C. briggsae*, which like *C. elegans* is hermaphroditic and supports RNA-mediated interference (Zhao *et al.* 2010). A third, and more important limitation, is that very few embryonic GRNs are known at high resolution in *C. elegans* that could serve as a comparison.

The specification of the *C. elegans* endoderm is an example of a set of interacting transcription factors that has been studied in great detail (Maduro 2017). In the early embryo, the founder cells E and MS are born (Fig. 1A). The E cell generates the entire endoderm (intestine), while its sister cell MS generates many mesodermal cell types, including the part of the pharynx, and many body muscle cells (Sulston *et al.* 1983). Many components of the GRN underlying MS and E development are known with high precision, and in most of cases, regulatory inputs have been confirmed to be direct and *cis-*regulatory sites have even been identified in upstream regions (Broitman-Maduro *et al.* 2006; Broitman-Maduro *et al.* 2005; Du *et al.* 2016; Maduro *et al.* 2001; Wiesenfahrt *et al.* 2015). This network is therefore a highly suitable system in which to examine questions of GRN evolution and developmental system drift.

**Fig. 1.**
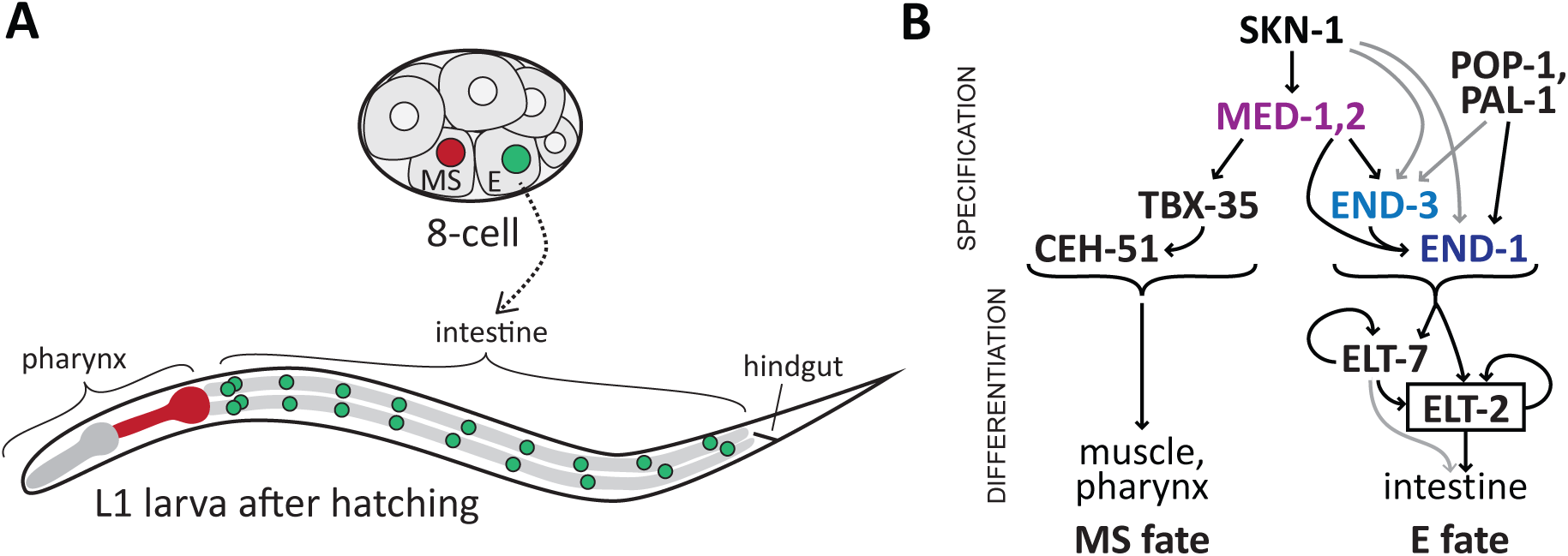
Embryonic origin of the E blastomere and simplified diagram of the gene regulatory network for endomesoderm specification in *C. elegans*. (A) The E cell and its sister cell MS are found ventrally in the 8-cell embryo (approximately 50 μm long). MS generates mesodermal cells including body muscles and the posterior portion of the pharynx, shown in red on the diagram of the larva (approximately 200 μm long). E generates the 20 cells of the intestine, whose nuclei are shown in green on the larva. (B) Specification of MS and E fates begins with the same SKN-1 and MED-1,2 factors, but then bifurcates into an MS pathway that includes the T-box factor TBX-35 and the homeobox factor CEH-51, while endoderm specification involves activation of END-3 and END-1. These upstream transient factors ultimately activate ELT-2 (and its paralogue ELT-7) which maintain intestinal fate. Additional input into E specification occurs by input from TCF/POP-1 and Caudal/PAL-1. All of MED-1,2, END-1,3 and ELT-2,7 are GATA type transcription factors.

The endomesoderm specification network works as follows. A simplified diagram is shown in Fig. 1B. Specification of both MS and E begins with accumulation of maternal SKN-1 protein. SKN-1 is an unusual transcription factor that binds DNA as a monomer through a Skn domain consisting of a homeodomain-like amino half recognizing an A/T-rich sequence, and a bZIP-like carboxyl basic domain recognizing a TCAT sequence (Blackwell *et al.* 1994; Carroll *et al.* 1997; Lo *et al.* 1998; Pal *et al.* 1997). SKN-1 directly activates expression of *med-1* and *med-2*, which encode nearly identical divergent GATA-type transcription factors that recognize an atypical AGTATAC core site (Broitman-Maduro *et al.* 2005; Lowry *et al.* 2009). SKN-1 and MED-1,2 are important for specification of both MS and E, as loss of activity of these genes results in a penetrant failure to specify MS, and an incompletely penetrant failure to specify E (Bowerman *et al.* 1992; Maduro *et al.* 2001). In MS, the MEDs specify mesodermal fate in part through activation of *tbx-35* (Broitman-Maduro *et al.* 2006). In E, SKN-1 and MED-1,2 contribute to activation of the paralogous *end-1* and *end-3* genes. These encode similar GATA factors that are expressed in the early E lineage, with *end-3* being activated slightly earlier than *end-1* (Baugh *et al.* 2003; Maduro *et al.* 2005a; Maduro *et al.* 2002; Zhu *et al.* 1997). In turn, the END-3 and END-1 proteins activate *elt-2*, a GATA factor that sets and maintains, through positive autoregulation, the fate of intestinal cells and is the central regulator for all intestinal genes (Fukushige *et al.* 1998; Fukushige *et al.* 1999; Mcghee *et al.* 2009). The *elt-7* gene encodes a similar GATA factor that shares function with *elt-2*, but which itself is not essential for normal development (Dineen *et al.* 2018; Sommermann *et al.* 2010). All of END-1, END-3, ELT-2 and ELT-7 seem to have similar DNA-binding properties and interact with canonical GATA binding sites of the type HGATAR (Du *et al.* 2016; Wiesenfahrt *et al.* 2015). Many additional studies have revealed unexpected nuance and complexity to the myriad of factors in this network, confirming that the sum of upstream inputs into *elt-2* activation is not merely additive. Upstream factors have distinguishable roles in establishment of robust cell divisions, gut morphogenesis and activation of genes important for metabolic function of the intestine (Boeck *et al.* 2011; Choi *et al.* 2017; Dineen *et al.* 2018; Maduro *et al.* 2015; Sawyer *et al.* 2011).

Integrated with the SKN-1 → MED-1,2 → END-1,3 feed-forward regulatory chain is the Wnt/β-catenin asymmetry pathway, which acts in the asymmetric MS *vs.* E fate decision through the nuclear effector TCF/POP-1 (Lin *et al.* 1995; Maduro *et al.* 2002; Owraghi *et al.* 2010; Rocheleau *et al.* 1997; Shetty *et al.* 2005; Thorpe *et al.* 1997). In MS, POP-1 represses gut fate by preventing activation of *end-1* and *end-3*, while in E, POP-1 is an activator that contributes to activation of *end-1* through its association with a divergent β-catenin, SYS-1 (Maduro *et al.* 2005b; Shetty *et al.* 2005). The POP-1 contribution to gut specification is not the major regulatory input, however, because loss of *pop-1* still results in endoderm specification from E (Lin *et al.* 1995). The contribution of POP-1 is detectable when depletion of *pop-1* is combined with loss of *skn-1*, *med-1,2* (together) or *end-3*, which produces loss of gut specification in a majority of embryos (Maduro *et al.* 2005a; Maduro *et al.* 2015; Maduro *et al.* 2007; Maduro *et al.* 2005b; Owraghi *et al.* 2010; Shetty *et al.* 2005). An additional minor input into gut specification in *C. elegans* is through maternally provided PAL-1 protein, a Caudal-like factor whose primary role is specification of a different blastomere called C (Hunter and Kenyon 1996; Maduro *et al.* 2005b).

A small number of studies have investigated the evolutionary dynamics of gut specification in species closely related to *C. elegans*. In *C. briggsae*, the *end-1* and *end-3* orthologues (the latter of which is found as two nearby paralogues, *end-3.1* and *end-3.2*) are expressed in the early E lineage, and knockdown of both by RNAi results in a failure to specify gut (Lin *et al.* 2009; Maduro *et al.* 2005a). In *C. briggsae* and *C. remanei*, most orthologues of the *med* genes, when introduced individually as transgenes, can fully complement the embryonic lethality of *C. elegans med-1,2(-)* embryos (Coroian *et al.* 2005). Together these studies suggested that the *med* and *end* factors play similar roles in all three species, as might be expected. Somewhat unexpectedly, however, knockdown of *skn-1* and *pop-1* orthologues in *C. briggsae* was found to produce different phenotypes from *C. elegans*, suggesting that the way that SKN-1 and POP-1 interact with their downstream target genes is subject to evolutionary changes even among very closely related species, i.e. the hallmark of developmental system drift (Lin *et al.* 2009; Zhao *et al.* 2010). From these few studies, then, a model emerges of a core endoderm specification pathway, where some regulatory inputs into the pathway are subject to more rapid evolutionary change than others.

An important way that properties of a GRN can be studied on an evolutionary scale is to examine features of orthologous genes in related species (Peter and Davidson 2016). However, given the essential requirement for the gut specification network in *C. elegans*, a paradox became apparent when genome sequences outside of the genus were completed: No *med* or *end* orthologues could be identified in the related nematode *Pristionchus pacificus*, while putative orthologues of *elt-2* and *skn-1* can be found in *Pristionchus* and in even more divergent species (data not shown) (Couthier *et al.* 2004; Dieterich *et al.* 2008; Schiffer *et al.* 2014). In recent years, however, the number of known species within the *Caenorhabditis* genus has grown considerably, opening possibilities for studying evolution of development through sequence comparisons (Kiontke *et al.* 2011). In the past two years, new sequence assemblies have become available for over two dozen *Caenorhabditis* genomes both within and outside of the so-called "Elegans supergroup" of species that are most closely related to *C. elegans* (Felix *et al.* 2014; Stevens *et al.* 2019). Collectively, this powerful set of sequences captures tens of millions of years of genome evolution (Cutter 2008; Stein *et al.* 2003).

In this work, I have taken a purely *in silico* approach and performed searches of *Caenorhabditis* genome sequence assemblies to identify orthologues of the *med*, *end-3* and *end-1* factors (Haag and Thomas 2015). Patterns of conservation of gene structure, protein structure and putative *cis*-regulatory sites are revealed in the *med* and *end* genes that confirm known information from *C. elegans* and reveal new insights into the MED and END proteins and the evolutionary dynamics of the network. The results complement studies that identify genome-wide conserved putative *cis-*regulatory motifs among close relatives of *C. elegans* (Grishkevich *et al.* 2011; Siepel *et al.* 2005; Zhao *et al.* 2012). A surprising finding is that the endoderm network likely originated at the base of the Elegans supergroup, in a manner that can be hypothesized to have resulted from the rapid serial intercalation of successive duplications of an ancestral GATA factor, likely *elt-2*. Other unexpected findings are the MED, END-3 and END-1 proteins are evolving at different rates, and that END-1 contains previously unrecognized, highly conserved domains that distinguish it from END-3. The resulting suite of MED/END-3/END-1 factors from 20 species forms a starting point for future studies on GRN evolution in *Caenorhabditis*.

## Materials and Methods

### IDENTIFICATION OF PUTATIVE MED AND END ORTHOLOGS

Sequence scaffolds and predicted proteins were downloaded from the *Caenorhabditis* Genomes Project (CGP) website (http://download.caenorhabditis.org) in late 2017. Searches were performed using the NCBI Windows 64-bit BLAST 2.7.1+ executable (ftp://ftp.ncbi.nlm.nih.gov/blast/executables/LATEST/) on a 64-bit Core i7 PC running Microsoft Windows 10, complemented by searching on both the CGP site and WormBase (http://wormbase.org). FASTA files containing sequence scaffolds, and others containing protein predictions, were searched by TBLASTN and BLASTP respectively using the protein sequences of *C. elegans* MED-1, END-1 and END-3. The updated *C. elegans* VC2010 sequence was also searched to confirm the *med* and *end* genes (Yoshimura *et al.* 2019).

Putative orthologous genes were identified using recommended best practices (Haag and Thomas 2015). Genes were first predicted by matching high-scoring segment pairs from TBLASTN results with genomic sequence, predicting the gene structure by identifying consensus intron splice donor and acceptor sequences, and comparing with the predicted genes from the assembly projects (Spieth *et al.* 2014; Stevens *et al.* 2019). Identification of gene structure started with the coding region for the DNA-binding domains and progressed both upstream and downstream. As analysis progressed, conserved features of the *med* and *end* genes and their gene products, within and among closely related species, became apparent, and these were used to refine the gene predictions. Searching of representative orthologs from each species back to the *C. elegans* genome confirmed that the predictions were the best matches. In some cases, the gene predictions from the assembly projects included short (<50 bp) predicted introns that could also be read through as coding. For these, a case-by-case judgment was made as to whether to include such introns in favor of maximizing amino-acid level homology. Some of the predictions within less-conserved regions could be incorrect, but these would not be expected to dramatically affect the analysis presented here. Similar judgments were made when multiple in-frame start codons were possible at the 5’ end of a gene, or when open reading frames could be extended in the 3’ direction by splicing around a stop codon. While no molecular validation of predicted genes was made, the manual curation of gene predictions favoring maximal similarity of gene and protein structures provides a surrogate validation by conservation across related species. This is the approach taken computationally for gene predictions by algorithms such as TWINSCAN (Korf *et al.* 2001).

It is highly likely that the gene set described here includes false duplicates. The quality and coverage of the genome assemblies, as well as the maintenance of heterozygosity in sequenced strains, are known to produce artifactual paralogues that are really alleles of one locus (Barriere *et al.* 2009; Haag and Thomas 2015). Some of these may still have been included as orthologues because they corresponded to a predicted gene from the sequence assembly. For example, the two *end-1* genes in *C. brenneri* are nearly identical with one found on a small sequence scaffold, suggesting that there is only one *end-1* orthologue in this species. The occurrence of these false duplicates is not expected to affect inter-species comparisons, for which a representative single gene/protein was chosen. Within a single species, a false duplicate would appear as a pair of nearly identical proteins. Gene models categorized as pseudogenes were more straightforward to find because they were truncated, had in-frame stop codons or frame shifts in the DNA-binding domain, or were missing essential amino acids such as one of the four cysteines in the C4 zinc finger. These may be expressed genes but were deemed unlikely to result in a functional protein.

Comparison of the protein predictions to the gene predictions of the various sequence projects validated the approach used to identify *med* and *end* orthologues. Of the genes identified and deemed not to be pseudogenes, 94/174 (54%) were identical to a predicted CDS from the assemblies, 56/174 (32%) partially overlapped an existing CDS, and 24/174 (14%) did not correspond to a predicted CDS. Differences from assembly project predictions often resulted from missing carboxyl and/or amino ends because of large introns, or extensions of open reading frames that maximized ORF length only. Completely missed predictions tended to be of the small intronless *med* genes that are often missed by gene-finding algorithms. Reliance of cDNA sequence data were not found to be useful, likely because the transient expression of the *med* and *end* factors in the earliest stages of embryogenesis meant that *med* and *end* RNAs were generally absent from mixed-stage cDNA preparations.

Predicted genes/proteins have been provisionally named *med-1.*n/MED-1.n, *end-3.*n/END-3.n, and *end-1.*n/END-1.n (where n = 1, 2, 3, etc.). Lower numbers correspond roughly to the rank order of identified high-scoring segment pairs from the TBLASTN search, which favors both stronger similarity with the *C. elegans* search sequence and scaffolds that contain multiple hits. Where a single orthologue was found in a species, it was named as *med-1*/MED-1, *end-1*/END-1 or *end-3*/END-3. For analyses where a single representative of a set of paralogues was used, it was the first numbered one, except for pseudogenes or one of the apparent two-fingered MEDs, in which case the next paralogue was used.

### IDENTIFICATION OF CONSERVED REGULATORY MOTIFS

A representative set of promoters, one per Elegans supergroup species per factor, was compiled to identify putative *cis*-regulatory motifs. This was done to reduce artifacts arising from overrepresentation of sets of very similar promoters resulting from intraspecific paralogs, which tended to have very similar promoters (data not shown). To identify sites starting with known binding sites, a JavaScript program was written to count occurrence of sites and compute p-values assuming a Poisson distribution, after the approach used in a prior work (Maduro *et al.* 2015). To identify motifs *ab initio* by their conservation, MEME (http://meme-suite.org/tools/meme) was used with expected site distribution with any number of repetitions (anr), the number of motifs to be identified as 10, and a maximum motif width of 12. Alternative parameters generally retrieved the same highly represented sites, except that motifs with higher E-values (and hence less conserved) could be different. Searches of the *end-1* and *end-3* promoters as separate groups produced qualitatively similar results as those that used both together, except that MED-like sites became rare enough among the *end-1* genes that they were not reported as significant by MEME. I did not consider sites whose E-values were greater than 1e-02 as these occurred among a small number of *med* and/or *end* genes. Some of these may represent less-conserved regulatory motifs, although they were not recognized as belonging to known factors from *C. elegans*. The site locations and promoter sequences are in Supplemental File S1.

### PHYLOGENETIC ANALYSIS

Alignments and simple Maximum-Likelihood trees were performed using MUSCLE as implemented in MEGA-X (Edgar 2004; Kumar *et al.* 2018). The tree for the DNA-binding domains was produced using RAxML as implemented in the RAxML-NG web service (https://raxml-ng.vital-it.ch) with default parameters, except that the BLOSUM62 substitution matrix was used and bootstrapping was activated (Kozlov *et al.* 2019; Stamatakis 2014). I note that construction of trees using the proteins described here results in disagreements with the more robust trees of Stevens *et al*. (2019), with only closely related species retaining the same relationship, such as the interfertile species *C. briggsae* and *C. nigoni* (Woodruff *et al.* 2010). This is what would be expected from rapidly evolving genes. Consistent with this, calculations of synonymous and non-synonymous substitutions rates did not produce interpretable information because of the high rates of molecular evolution in *Caenorhabditis* in general (Cutter 2008). Moreover, the fastest rates of evolution in *Caenorhabditis* occur in early zygotic regulators with transient expression, which accurately describes the MED and END factors (Cutter *et al.* 2019). Because fast-evolving proteins are being compared among 20 species (as opposed to only two or three), the major conclusions regarding conserved amino acids and stringency of selection are nonetheless self-evident from the alignments and shape of phylogenetic trees.

### ADDITIONAL SOFTWARE

Gene modeling, sequence alignments and other analyses were performed with Vector NTI 6 and the MEGA-X software package (Kumar *et al.* 2018). Generation of tables and drawing of to-scale diagrams in SVG format were aided by custom programs written by the author in JavaScript and Python. These scripts are available by request. Protein alignments were annotated using BoxShade (https://embnet.vital-it.ch/software/BOX_form.html) to generate EPS-formatted files. Data were compiled in Microsoft Excel and figures were assembled in Adobe Illustrator.

### DATA AVAILABILITY

Sequences identified in this work are available as Supplemental Files through **figshare** under “Maduro,2019-SupplementalFiles.”

## Results

### MED, END-3 AND END-1 ARE FOUND TOGETHER IN THE ELEGANS SUPERGROUP SPECIES

I searched sequence scaffolds from 27 species of the *Caenorhabditis* Genomes Project (http://caenorhabditis.org) with TBLASTN using the protein sequences of *C. elegans* MED-1, END-3 and END-1. *C. elegans*, *C. briggsae* and *C. remanei* were included as their sequences have been updated since earlier reports on *med* and *end* genes from these (Coroian *et al.* 2005; Maduro *et al.* 2005a; Yoshimura *et al.* 2019). As shown in Fig. 2, at least one orthologue of each of the three genes was found in 20 species comprising the Elegans supergroup, a clade that includes the Japonica and Elegans groups (Kiontke *et al.* 2011; Stevens *et al.* 2019). Consistent with the absence of even more distant MED or END orthologues, the number of putative GATA factors in the genomes of species outside the Elegans supergroup was smaller, typically 5 or fewer, and putative orthologues were better matched to other *C. elegans* GATA factors like ELT-3 (data not shown). Across the 20 species searched in the Elegans supergroup, *end-1* orthologs were unique in each genome except for *C. brenneri* (which has two *end-1* genes), while multiple paralogs within a species was the norm for the *end-3* orthologs with an average of 2.0 times per genome, and the *med* orthologues, found an average of 5.6 times. Of 208 genes identified, 34 were deemed to be the result of unresolved heterozygosity or were likely pseudogenes (counted together under "pseudo" in Fig. 2); these were eliminated from further study. It is still likely that some false duplicates persist in the predicted gene set, so occurrence of nearly identical paralogues should be interpreted with caution (see **Materials and Methods**). In any event, the identification of false duplicates would not change the results of inter-species comparisons, for which a single representative gene was chosen for each factor.

**Fig. 2.**
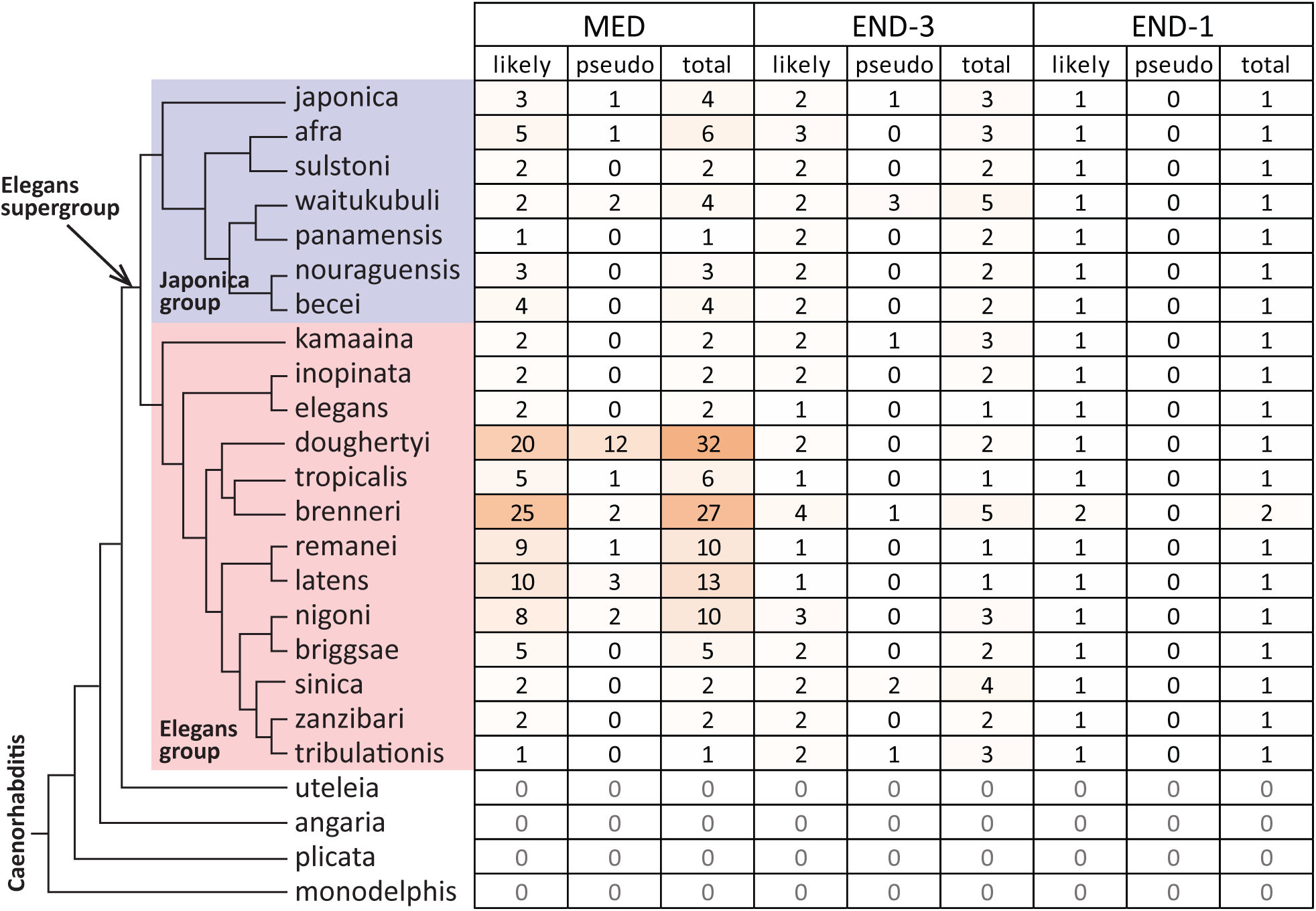
Orthologues of the MED, END-3 and END-1 genes among species whose sequences were searched. Species are shown after the phylogeny in (Stevens *et al.* 2019) with the Japonica group in light blue and the Elegans group in pink. The species *C. parvicauda, C. castelli, C. quiockensis*, and *C. virilis*, which contained no orthologues of the MED and END factors, have been omitted for simplicity. Table cells are colored by the number of orthologues.

### CONSERVED LINKAGE OF *end-1* and *end-3* ORTHOLOGUES

In *C. elegans* and *C. briggsae* the *end-1* and *end-3* genes are within ~30 kbp of each other (Maduro *et al.* 2005a). Microsynteny of this type has been observed in other genes of these two species (Coghlan and Wolfe 2002; Kent and Zahler 2000). To see if microsynteny of *end-1* and *end-3* is common, I examined whether *end-1* and *end-3* orthologues in other species may be linked. As shown in Fig. 3A, in 12/18 of the remaining Elegans supergroup species, *end-1* and *end-3* are found on the same scaffold with an average separation of ~37 kbp and a range of 20-63 kbp. In *C. brenneri*, which has two *end-1* and five *end-3* orthologues, one scaffold carries both an *end-1* and an *end-3*, however the distance between them is ~530 kbp. In the remaining five species, the *end-1* and *end-3* genes are found on different scaffolds. Because it is possible for a sequence scaffold to break between two linked genes, there may be additional synteny among these. For example, in *C. sinica* the scaffold containing the *end-1* orthologue is 32 kbp in size with the *end-1* gene located 3 kbp from one end, raising the possibility that although its *end-3* ortholog is on a different scaffold, *end-1* and *end-3* may be nearby in the genome. Closely related species have similar patterns of *end-1* and *end-3* synteny, for example between *C. afra* and *C. sulstoni*, and between *C. zanzibari* and *C. tribulationis* (Fig. 3A). Although synteny is conserved, the relative orientation of linked *end-1* and *end-3* paralogues varies, with examples of all four possible linked arrangements. In *C. elegans*, *end-1* and *end-3* are encoded on the same strand with *end-1* upstream of *end-3*. In *C. sulstoni*, two *end-3* paralogs are upstream of *end-1* with all three genes on the same strand. In *C. zanzibari* and *C. tribulationis*, *end-1* is on one strand in between two *end-3* paralogs on the other strand, hence in one *end-1/3* pair the genes point towards each other, and in the other they are divergently transcribed. These differing arrangements are consistent with the high rate of intrachromosomal rearrangements previously noted for *Caenorhabditis* (Coghlan and Wolfe 2002).

**Fig. 3.**
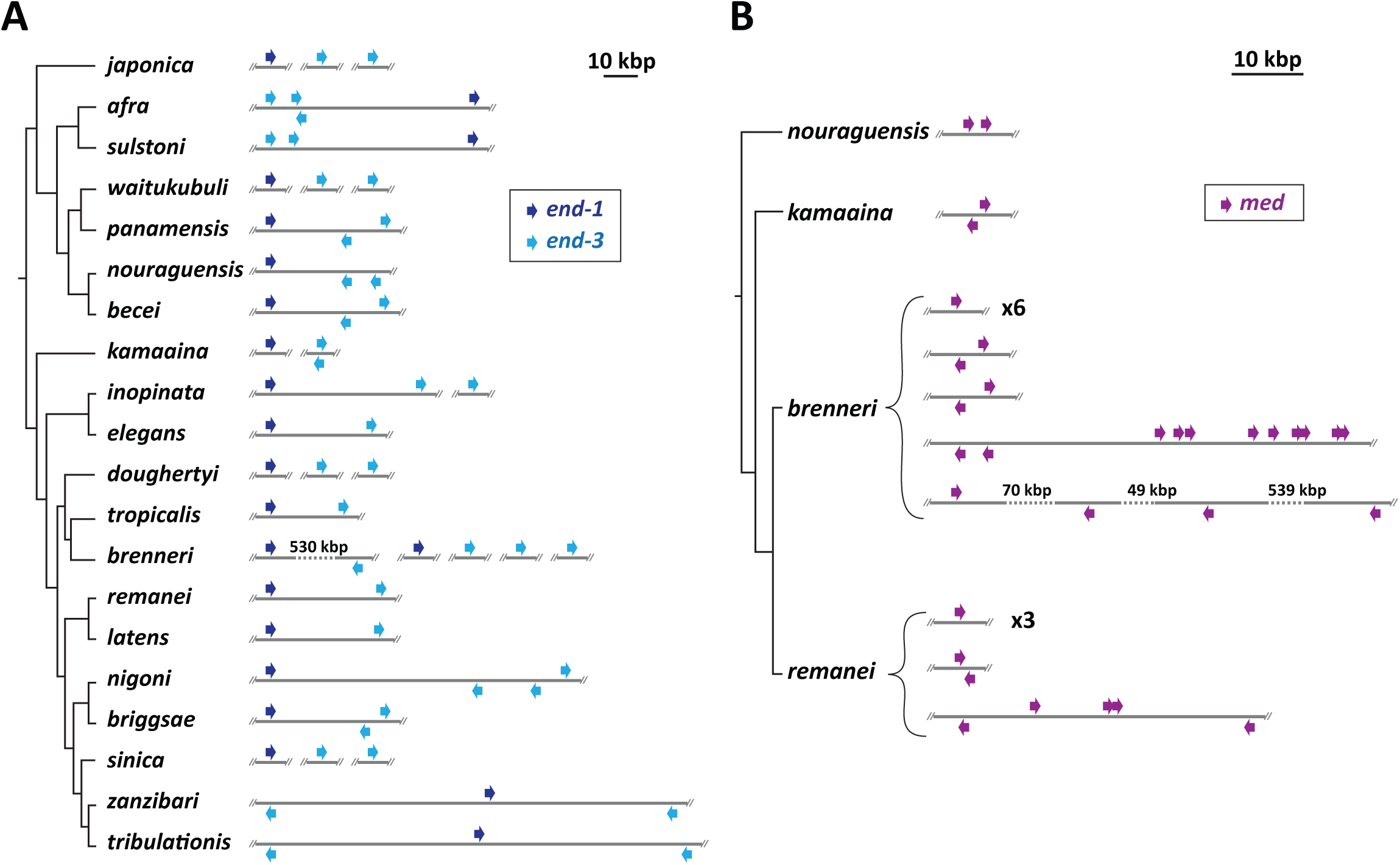
Synteny and relative orientation among *med* and *end* genes found on sequence scaffolds. Except where noted by a number, inter-gene distances are shown relative to the scale bar at the top of each panel. (A) Patterns of linkage among *end-1* (dark blue) and *end-3* (light blue) orthologues among the Elegans supergroup species. (B) Patterns of linkage among *med* orthologues for a subset of species.

### PREVALENCE OF LINKED *med* AND *end-3* DUPLICATIONS

In *C. briggsae*, two *end-3* paralogues are found in an inverted orientation within several kbp, and in *C. remanei*, two clusters of closely linked *med* paralogues were found (Coroian *et al.* 2005; Maduro *et al.* 2005a). Similar linked duplications of these genes were found in other species. Among the *end* genes shown in Fig. 3A, 7/10 species with at least two *end-3* genes show two of them within 10 kbp. Among the 18 species with at least two *med* genes, linked pairs can be found in nine of them, in which at least two *med* genes occur within 5 kbp of each other. Examples of linked *med* duplications are shown for four of the Elegans supergroup species in Fig. 3B. In the most extreme case, 9/25 *C. brenneri med* orthologs are clustered across a 23-kbp region, with an additional tandem pair located ~22 kbp away. Linked duplications are therefore a common occurrence, particularly for the *med* genes.

### ABSENCE OF A CONSERVED INTRON IN THE ELEGANS GROUP *med* GENES

I next examined the evolutionary changes in *med* and *end* gene structures across the Elegans supergroup. For simplicity, a single representative *med*, *end-3* and *end-1* gene was used for each species because intraspecific paralogs generally showed identical splicing patterns. The gene structures are shown in scale diagrams in Fig. 4A, depicting intron/exon structures arranged by the phylogeny of Stevens et al. (2019). Intron positions are also indicated on diagrams of the predicted proteins in Fig. 8. Of particular significance, prior work found that the *med* genes of *C. elegans*, *C. briggsae*, and *C. remanei* have no introns, unlike all other GATA factors in these species including the *end* genes (Coroian *et al.* 2005; Gillis *et al.* 2008; Maduro *et al.* 2001). As shown in Fig. 4A, while all representative *med* genes were found to be intronless across the Elegans group, the *med*s from the Japonica group share a common intron (indicated by an asterisk) within the C4 zinc finger coding region that is found in the same position in all *end-1* and *end-3* genes. In addition to this conserved intron, within the Japonica group, the *C. japonica* and *C. panamensis med* genes each have one more upstream intron at non-homologous positions.

**Fig. 4.**
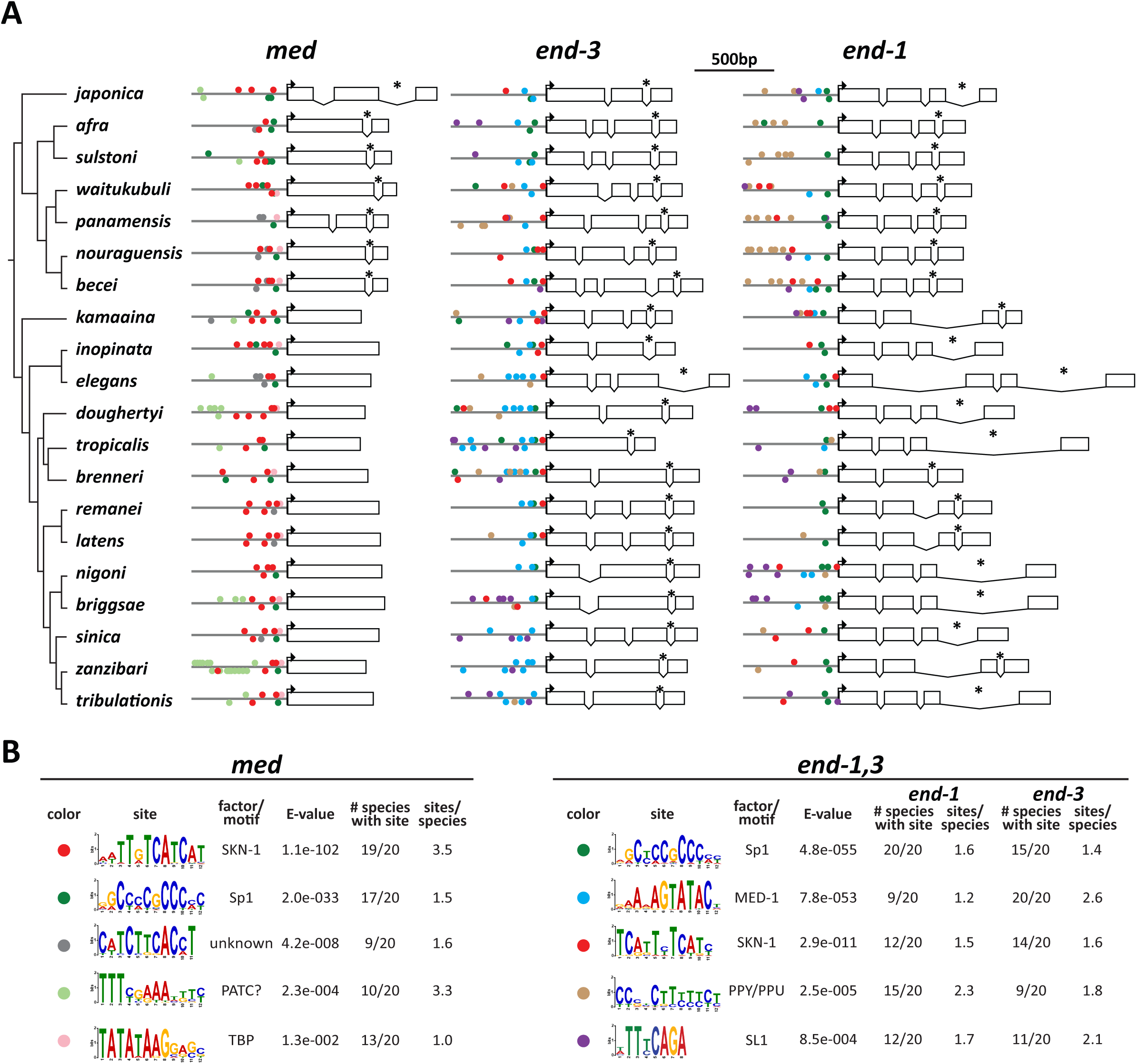
*med* and *end* gene structures and conserved promoter motifs. (A) Gene structures. 600bp of promoter are shown as a line, and the coding DNA sequence (CDS) predictions are shown relative to the scale bar at the top. Boxes are exons, and spaces joined by a ‘V’ are introns. Bent arrows indicate the location of the predicted start codon. An asterisk denotes the intron conserved among all *end* genes and Japonica group *med* genes. (B) Motifs identified by MEME for the *med* and *end-1,3* genes. The motifs are symbolized by a colored circle on the promoters in (A). Some of the motifs are shown in their reverse complement from the MEME output files in Supplemental Files S13 and S14.

### DIFFERENCES IN INTRONS AMONG *end-3* AND *end-1* GENES

The conserved zinc finger intron is the only one shared between the *end-3* and *end-1* genes (Fig. 4A). As a group, the *end-3* orthologs show the highest variability in the number of introns, with *C. tropicalis* having only the one conserved intron, *C. becei* having four introns total, and the remaining species having two or three. The *end-1* orthologues are far less diverse, sharing the same four exons with three introns, except for *C. brenneri* which is missing the second intron. In terms of size, the *end-3* introns tend to be smaller overall, with introns larger than 100 bp most apparent within the Elegans group *end-1* genes. Hence, the positions of introns in the *end-1* orthologues appear to be under a greater constraint than those of the *end-3* genes.

### IDENTIFICATION OF CONSERVED PROMOTER MOTIFS

The occurrence of *med* and *end* genes in 20 related species affords the opportunity to identify conserved *cis-*regulatory sites and infer conservation of the structure of the gut specification network. The expectation is that conserved regulatory inputs found in *C. elegans* should be reflected in the occurrence of similar *cis*-regulatory sites mediating the same promoter-DNA interactions in the other species. I first searched for known binding sites for *C. elegans* factors among the Elegans supergroup *med* and *end* orthologues using methods previously used in *C. elegans* (Maduro *et al.* 2015). A size of 600bp upstream of the ATG was chosen for these and subsequent analyses, as the known regulatory interactions with the *C. elegans med* and *end* genes generally occur within a few hundred base pairs of the ATG (Bhambhani *et al.* 2014; Broitman-Maduro *et al.* 2005; Maduro *et al.* 2001; Shetty *et al.* 2005). Among the *med* upstream regions, I found only widespread conservation of SKN-1-like sites, and among the *end-3* orthologues, only MED sites (Supplemental Tables S1, S2 and S3). While these results support conservation of activation of *med* orthologues by a SKN-1-like factor, and activation of *end-3* orthologs by MED-like factors, a complementary (and superior) approach is to search for over-represented motifs *ab initio*. I therefore searched 600bp upstream of representative *med* and *end* genes from all 20 species using the MEME discovery algorithm (Bailey and Elkan 1994). The results are summarized in Fig. 4B, with the sites indicated by color coded circles on the promoters in Fig. 4A. The locations of the sites diagrammed in Fig. 4 are listed in Supplemental File S1.

### SKN-1 BINDING SITES IN THE *med* AND *end* GENES

Among the *med* orthologues, a motif resembling two overlapping SKN-1 sites was identified 19/20 species. The core of this motif, RTCATCAT, was found in two clusters in the *C. elegans med* genes and DNA fragments containing these sites are capable of binding recombinant SKN-1 DNA-binding domain *in vitro* (Maduro *et al.* 2001). The same core is found in SKN-1 binding sites in *gcs-1*, a known SKN-1 target gene in the fully developed intestine (An and Blackwell 2003). As in *C. elegans*, the SKN-1 sites in the *med* genes are found within 300 bp of the predicted start site in most of the other species, which is apparent from the diagram in Fig. 4A. In *C. panamensis*, which contains only a single putative *med* gene, an RTCATCAT site was not identified by MEME although six ‘core’ RTCAT sites were found by direct searching (p ≤ 0.05, Poission distribution). The low E-value of 1.1e-102 and presence of an average of 3.5 sites per species strongly suggest that activation of *med* orthologous genes likely occurs by SKN-1 in most Elegans supergroup species.

Among the *end-1* and *end-3* genes, a TC<u>ATT</u>Y<u>TCATC</u> site was identified by MEME in 12/20 *end-1* genes and 14/20 *end-3* genes (E-value 2.9e-11). Most of this site (underlined) overlaps with 8/9 bases of the WWWRTCATC site for SKN-1 (Etheve *et al.* 2016; Mathelier *et al.* 2014). Unlike the SKN-1 sites in the *med* genes, which occur an average of 3.5 times per gene, these putative SKN-1 sites in the *end* genes, when present, occur only 1.5 times per *end-1* gene and 1.6 times per *end-3* gene. I hypothesize that this site represents a degenerate SKN-1 binding site. Prior evidence in *C. elegans* had suggested that SKN-1 contributes directly to *end-1,3* activation independently of the MEDs, though the precise sites have not been reported (Maduro *et al.* 2015).

### Sp1 BINDING SITES

A motif resembling the binding site for Sp1 was found in the *med* promoters (17/20 species, E-value of 2.0e-33), *end-1* (20/20 species), and *end-3* promoters (15/20 species), with an E-value of 4.8e-55 for the two *end* genes. This same motif has been found among many *C. elegans* promoters, suggesting that regulation by Sp1 is not restricted to gut specification (Grishkevich *et al.* 2011). Reduction of function of *sptf-3*, a gene encoding an Sp1-like factor, causes a decrease in specification of E and a reduction in expression of *end-1* and *end-3* reporters (Sullivan-Brown *et al.* 2016). From the widespread conservation of the Sp1 binding sites, it is likely that Sp1 contributes to E specification across many species in the Elegans supergroup through direct binding of the *med*, *end-1* and *end-3* orthologous genes.

### MED BINDING SITES IN THE *end-1* AND *end-3* GENES

Prior work identified the binding sites for the MED factors in the *end-1* and *end-3* genes, defining a core sequence of AGTATAC that is distinct from the HGATAR site of canonical GATA factors (Broitman-Maduro *et al.* 2006; Broitman-Maduro *et al.* 2005; Lowry *et al.* 2009). As anticipated by the results from searching for this site directly, MEME identified a highly conserved MED site motif in 9/20 *end-1* genes and 20/20 *end-3* genes (E-value 7.8e-53 across both *end-1* and *end-3*). Across the nine species with MED sites identified in *end-1*, there are an average of 1.2 sites per gene, while for *end-3*, there are 2.6 sites on average. The location and spacing of the sites are consistent with results from *C. elegans*, with sites occurring within 200 bp of the predicted translation start site and showing a spacing (when multiple sites are present) of ~50 bp (Broitman-Maduro *et al.* 2005).

### POLYPYRIMIDINE MOTIF

MEME identified a pyrimidine-rich motif in 15/20 *end-1* genes and 9/20 *end-3* genes (E-value 2.5e-05). This motif, consisting primarily of C and T, is most apparent among the Japonica group *end-1* genes. The complement of the pyrimidine-rich motif is purine-rich, hence these motifs are called PPY/PPU (polypyrimidine/polypurine) tracts (Sawicka *et al.* 2008). This motif did show a strand bias by gene: 30/34 sites among the *end-1* genes have the polypyrimidines on the top strand, while the sites are evenly on either strand (9/16 on the top strand) in the *end-3* genes. Polypyrimidine tracts are generally associated with messenger RNAs where they would be present as one strand, and interact with polypyrimidine-tract binding proteins (PTBs) (Sawicka *et al.* 2008). Curiously, Pur-alpha-like protein (PLP-1), a factor that binds a purine-rich sequence, was previously identified as having a regulatory input into *end-1* activation in *C. elegans* (Witze *et al.* 2009). However, the PPY/PPU motif identified by MEME was not found in either of the *C. elegans end* genes.

### ADDITIONAL OVERREPRESENTED MOTIFS

Three additional sites were found by MEME among the *med* genes. A motif containing a TCTKCAC core was found in 9/20 species *med* genes with an average of 1.6 sites per gene (E-value 4.2e-08). The motif sequence does not immediately suggest a putative regulatory factor, although it tends to be found among the SKN-1 sites, suggesting it is related to SKN-1 binding. A motif containing TTTNNAAA was found at a higher E-value of 2.3e-04 in 10/20 *med* genes with an occurrence of 3.3 sites per gene, with one species *C. zanzibari*, containing 16 of them. This site resembles previously identified periodic AT clusters (PATCs) suggesting it may be a more general motif (Frokjaer-Jensen *et al.* 2016). A motif resembling a TATA-box was found in 13/20 species’ *med* genes with an even higher E-value of 1.3e-02 (Grishkevich *et al.* 2011). This may be a *bona fide* basal promoter site, as it is found within tens of base pairs from the translation start in these 13 genes. Finally, among the *end* genes, an "SL1 motif" was found in 12/20 *end-1* genes and 11/20 *end-3* genes (E-value 8.5e-04) (Grishkevich *et al.* 2011). The motif was not found in the *C. elegans end-1/3* genes, consistent with prior work that neither of these in *C. elegans* are not known to be *trans-*spliced to the SL1 sequence (Allen *et al.* 2011; Zhu *et al.* 1997). Its relevance as a motif is uncertain, as in most of the *end* promoters that contain it, the site is more than 300bp upstream of the predicted start site.

### PHYLOGENETIC ANALYSIS CONFIRMS THAT MED, END-3 AND END-1 FORM DISTINCT CLADES

The gene structure and promoter motifs suggest that the *med*, *end-3* and *end-1* genes form distinct families among the 20 species of the Elegans supergroup. To confirm that this is reflected at the protein level, I aligned the DNA-binding domains (DBDs) among representative MED, END-3 and END-1 factors (one per species) and used this to construct a phylogenetic tree *ab initio* with the RAxML-NG method (Kozlov *et al.* 2019; Stamatakis 2014). As shown in Fig. 5, MED, END-3 and END-1 form three broad clades, with the END-1 factors showing the highest similarity as a group, followed by the END-3 factors, and finally the more diverse MED factors. A high diversity of the MED factors was previously observed among the *med* genes from *C. elegans*, *C. briggsae* and *C. remanei* (Coroian *et al.* 2005). The grouping of the factors increases confidence that the correct orthologues have been assigned and shows that different rates of protein evolution have occurred among the three factors.

**Fig. 5.**
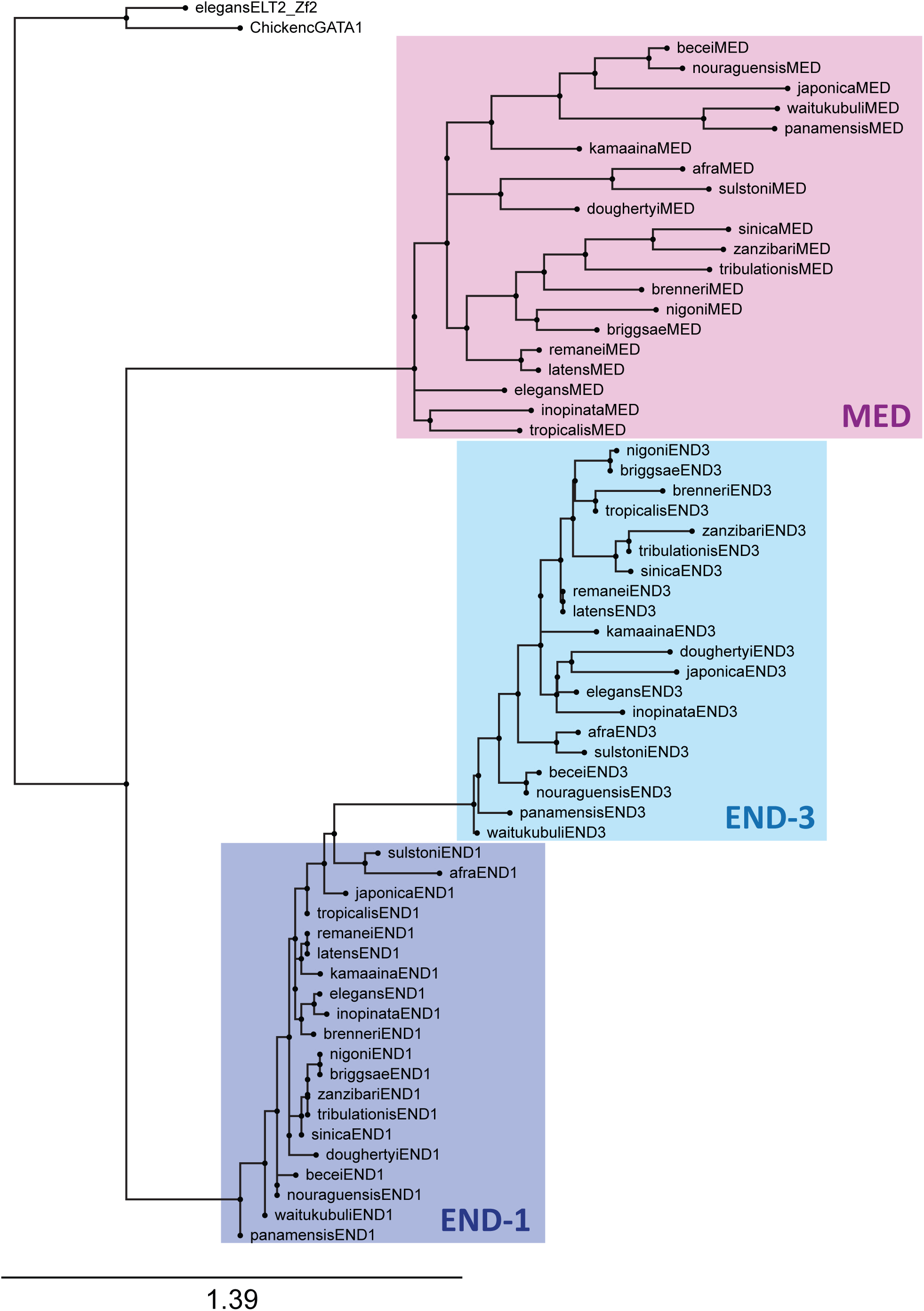
Phylogenetic tree of representative MED, END-3 and END-1 DNA-binding domains. The DNA-binding domains of *C. elegans* ELT-2 and chicken GATA1 are shown as outgroups. Each of the three factors forms a distinct clade, with the END-1 factors showing the highest similarity, followed by END-3, then the MEDs as the most diverse group.

### GENE AMPLIFICATION WITHIN AND AMONG SPECIES

While *end-1* is represented by a unique orthologue among all species (except *C. brenneri* which may have two *end-1* genes), *med* and *end-3* orthologues are often found as two or more duplicate genes within a species. The two *C. briggsae* END-3 paralogues are highly similar, suggesting recent duplication, and the multiple *med* genes among *C. elegans*, *C. briggsae* and *C. remanei* are also much more alike within each species (Coroian *et al.* 2005; Maduro *et al.* 2005a). To test how general this phenomenon is, I aligned and constructed trees for all MED DBDs, and separately, the END DBDs. In the tree of MED factors shown in Fig. 6, most *med* duplications have occurred post-speciation from a small number of founding genes. The 20 MED factors in *C. doughertyi* cluster in a way that suggests there may have been only one or two ancestral *med* genes that underwent multiple rounds of amplification. In the case of *C. brenneri*, the MEDs form two clusters of 22 and 3 genes each, suggesting there were only a few ancestral factors. A similar division occurs among the *C. tropicalis* MEDs, which suggests two ancestral *med* genes. There are three groups in which paralogous MED factors are clustered within species pairs: *C. briggsae* with *C. nigoni*, *C. becei* with *C. nouraguensis*, and *C. latens* with *C. remanei*. Within each cluster, the pattern suggests that both species inherited two or three *med* paralogues from a common ancestor, which then each underwent further amplification post-speciation. Among the remaining 9 species that have 2-5 *med* genes each, the paralogous MEDs clustered together as a single group, suggesting a single ancestral gene. This unusually widespread pattern of duplications both pre- and post-speciation, not seen in the *end* genes, shows that the *med* genes are under different evolutionary constraints.

**Fig. 6.**
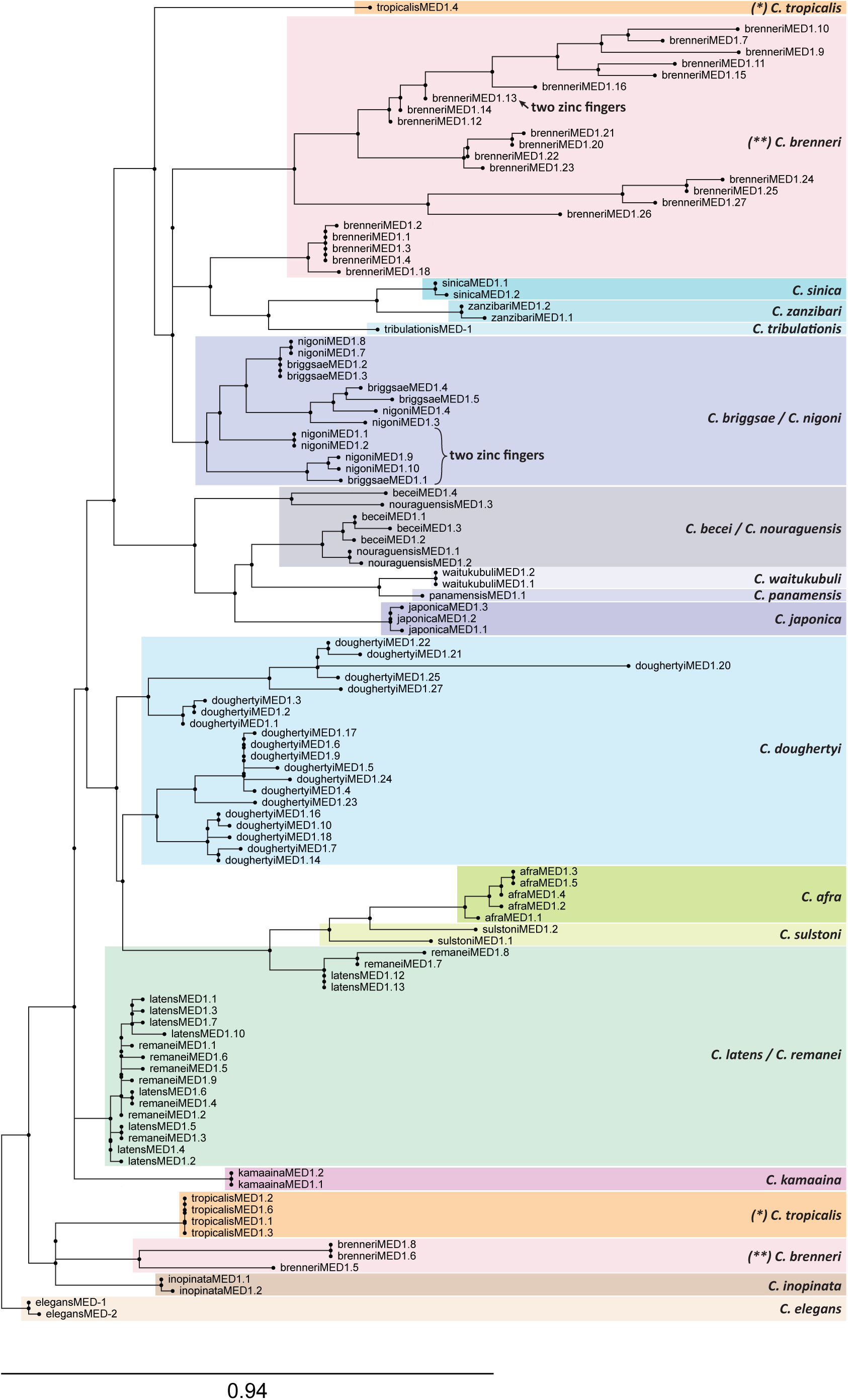
Phylogenetic tree of all MED factors, showing high prevalence of duplications across the Elegans supergroup. In most cases, paralogous duplicates likely arose post-speciation, although there are examples that suggest that some species each inherited two or three genes from a common ancestor that later underwent further duplications. The tree was generated by RAxML using the MED DNA-binding domains (Kozlov *et al.* 2019; Stamatakis 2014).

I note here that six genes were found that encoded MED-like factors with two C4 zinc fingers, indicated on the tree in Fig. 6. In each case, the two fingers were highly similar, so only one of the two fingers was used to generate the tree. Four of the genes were present as two paralogous pairs in *C. nigoni*, one was found in *C. briggsae*, and another was found in *C. brenneri* (Fig. 6). *C. nigoni* and *C. briggsae* are very closely related, suggesting they inherited the same two-fingered *med* gene from a common ancestor (Kiontke *et al.* 2011). The positions of the six two-fingered MED factors in the phylogeny are hence consistent with two-finger MED-type GATA factors having arisen twice, likely by an interstitial duplication, because the two fingers in each share a nearly identical amino acid sequence. The observation of putative two-fingered GATA factors is noteworthy because among vertebrates, GATA factors generally have two zinc fingers (Gillis *et al.* 2009; Lowry and Atchley 2000).

A tree of the DBDs of the END-1 and END-3 orthologues is shown in Fig. 7. As mentioned earlier, all END-1 orthologues are unique in each species except for the two possible *end-1* paralogues in *C. brenneri*. Among the END-3s, intraspecific amplification was implied for all species with two or more END-3s, except for a cluster containing END-3 paralogues from *C. sinica*, *C. tribulationis*, and *C. zanzibari*. This portion of the tree is most consistent with two paralogous *end-3* genes having been present in the common ancestor of all three species. Hence, duplications do occur among the *end-3* paralogues, but at a far lower frequency than with the *med* genes.

**Fig. 7.**
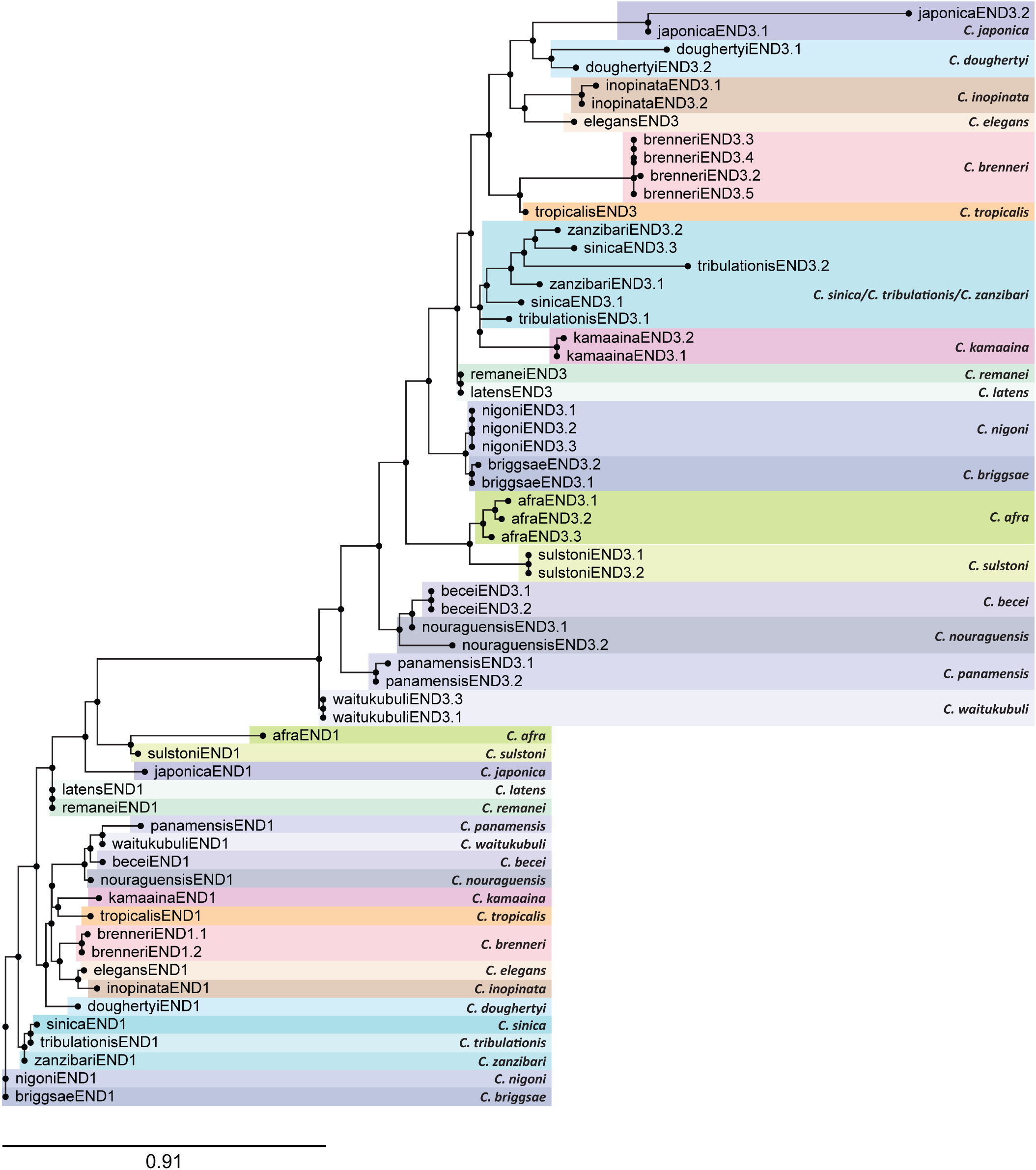
Phylogenetic tree of all END-3 and END-1 factors, showing tendency for END-1 factors to be unique, and END-3 factors to have undergone some duplications. The tree was generated by RAxML using the END-3 and END-1 DNA-binding domains (Kozlov *et al.* 2019; Stamatakis 2014).

### CONSERVED DOMAINS OF MED, END-3 AND END-1

Prior alignments of the ENDs from *C. elegans* and *C. briggsae* revealed three conserved domains: An amino-terminal polyserine (Poly-S) region, a short region immediately upstream of the zinc finger, called the Endodermal GATA Domain (EGD), and the GATA-type zinc finger and basic domains (Maduro *et al.* 2005a). Among the MEDs, only the latter two domains were conserved (Coroian *et al.* 2005). Taking advantage of the 20 Elegans supergroup species, we aligned representative MED and END proteins to both generalize these earlier findings and to identify other conserved domains that might have been missed. The alignments revealed both expected and previously unknown conserved regions, shown diagrammatically in Fig. 8. On this figure, the corresponding positions of introns are also indicated to reveal patterns of conservation of the gene structure in relation to these conserved regions.

**Fig. 8.**
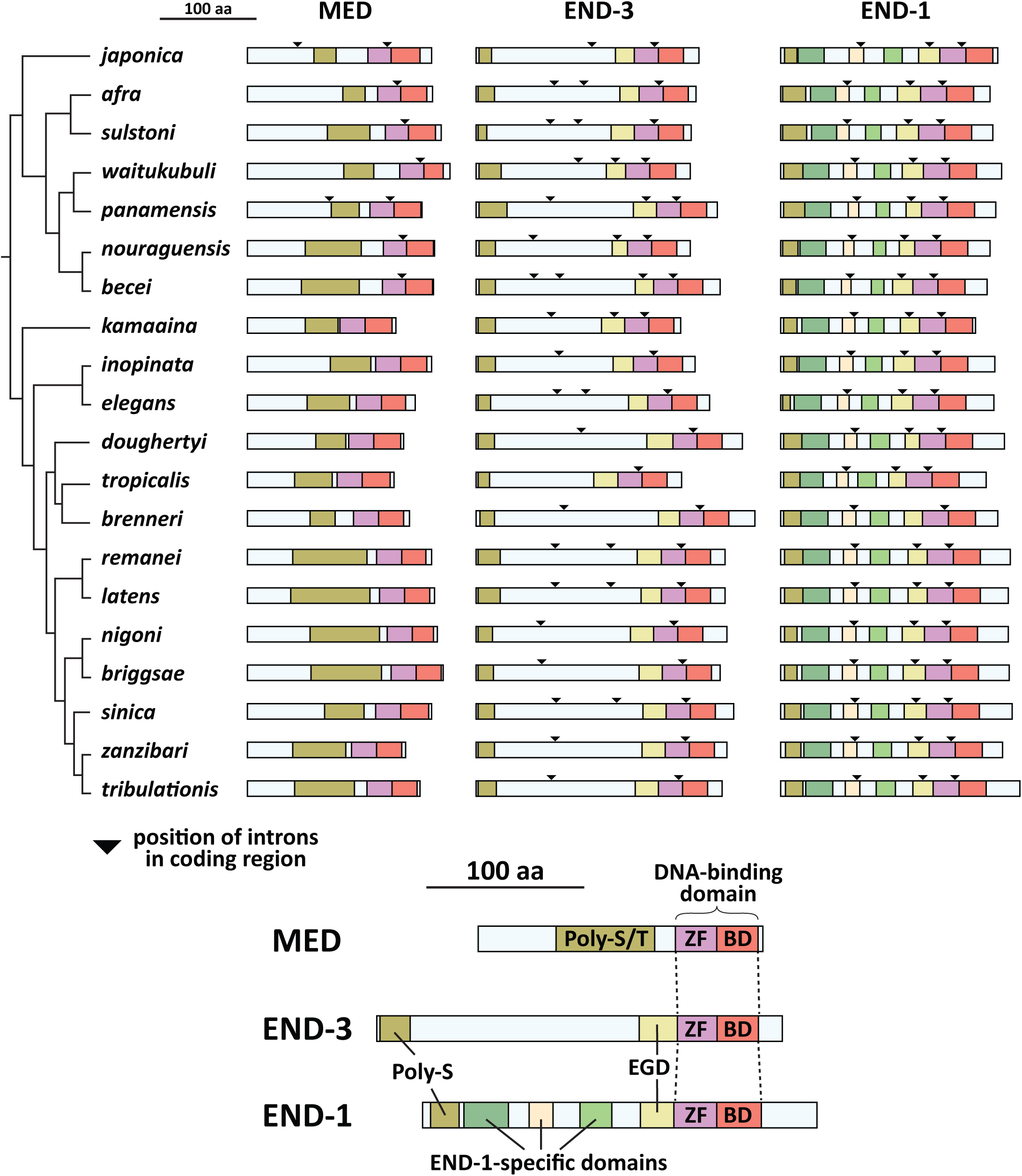
Conserved MED and END protein domains. The top part of the figure shows the MED, END-3 and END-1 protein structures with conserved domains in colored regions. Triangles represent the positions of introns in the coding regions as shown in the gene models in Fig. 4A. The bottom of the figure shows the names of the domains, which are shown at the amino acid level in Figs. 9 and 10. The MED orthologues have a variable region high in serine and threonine (Poly-S/T), while END-1 and END-3 share an amino-terminal polyserine domain (Poly-S) of variable length and an Endodermal GATA Domain (EGD). The END-1 orthologs share three additional regions not found in END-3. The species are arranged after the phylogeny in (Stevens *et al.* 2019).

### MED, END-3 AND END-1 DNA-BINDING DOMAINS

An alignment of representative DBDs for the MED, END-3 and END-1 factors, one per species, is shown in Fig. 9 (Edgar 2004). Consistent with their recognizing an atypical binding site, the MED DBDs share features that distinguish them from the END-3 and END-1 DBDs (Fig. 9A). Among the Elegans group MED factors, the C4 zinc finger has 18 amino acids between the two pairs of cysteines, with a structure of CXXC-X_18_-CXXC, while the Japonica group members are diverged from this structure and have 16-17 amino acids, i.e. CXXC-X_16-17_-CXXC. A consensus sequence with 11 invariant amino acids is shown below the alignment in Fig. 9A. While the group of MED factor DBDs appear to be diverse, the identification of a conserved MED-like motif among the *end-3* promoters suggests that the MED factors have nonetheless coevolved to continue recognizing a similar binding site in each species. The solution structure of a *C. elegans* MED-1 DBD::binding site complex revealed that recognition of the MED binding site is mediated by 9 amino acids, indicated at the bottom of Fig. 9A (Lowry *et al.* 2009). In comparing these with the corresponding amino acids in the other MED DBDs, there is evidence of conservation as shown by asterisks. Two of the 9 amino acids, a tyrosine (Y) and arginine (R) just after the zinc finger, are invariant. Five of the remaining amino acids are found in most of the MED DBDs. The remaining two are the isoleucine (I) and the first arginine in the zinc finger. The arginine is somewhat conserved, as in most MEDs it is an arginine or a lysine (K), both of which are basic. The isoleucine (I) is not conserved, however, and is replaced by a cysteine (C) in most other MEDs. This amino acid may not be critical for recognition of a MED binding site, however, as prior work showed that transgenes containing individual *med* genes from *C. briggsae* and *C. remanei* can fully complement the embryonic lethal phenotype of *C. elegans med-1; med-2* double mutants; in the MED factors from both of these species, the corresponding amino acid is a cysteine. Overall, despite the higher divergence among the MEDs as a group, there appears to be selection for the 8/9 amino acids known to be involved in site recognition in *C. elegans* MED-1. Added to the apparent conservation of MED-like binding sites in the respective *end-3* orthologues in every species, the data suggest maintenance of the DNA-binding specificity of the MEDs.

**Fig. 9.**
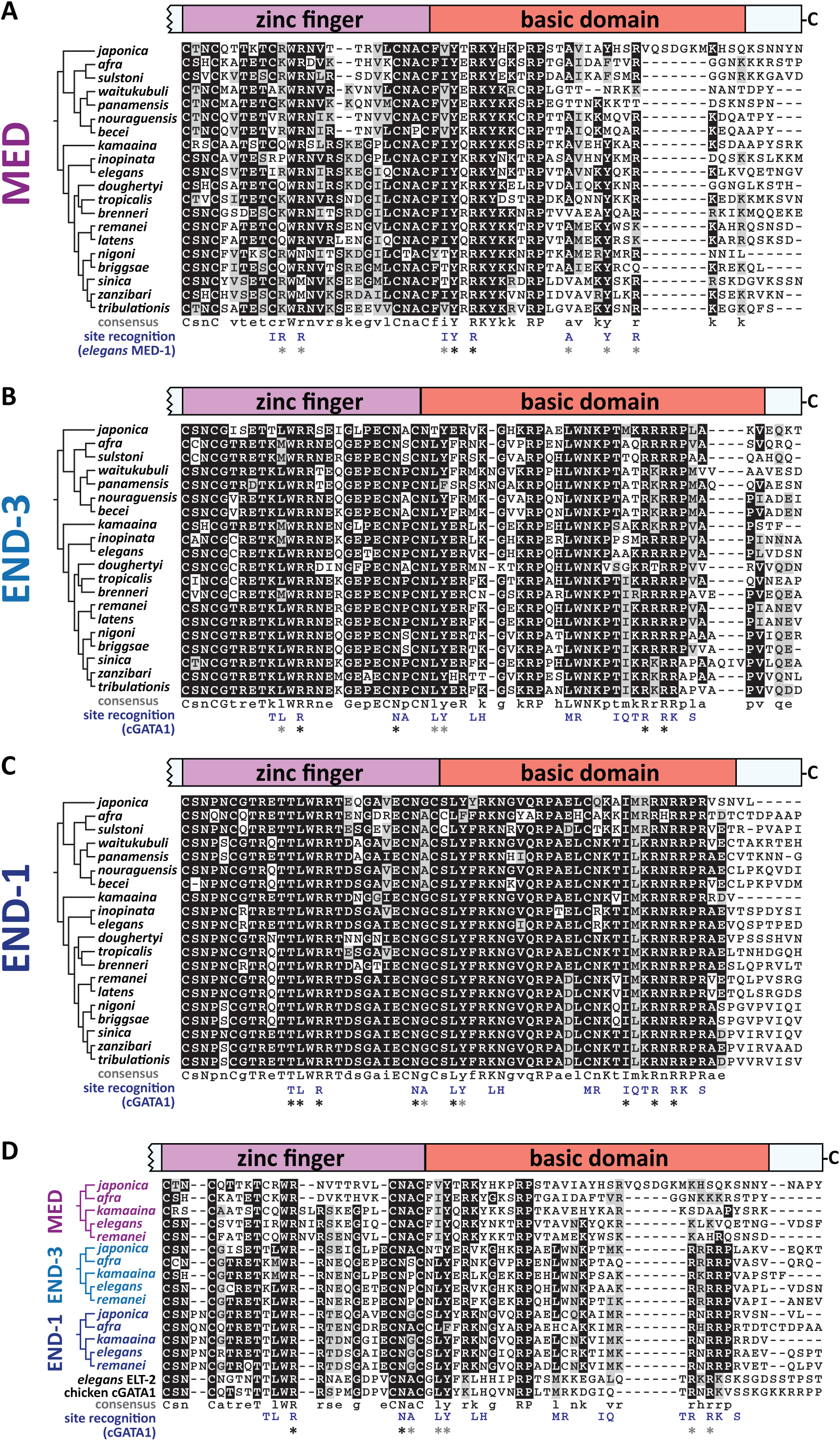
DNA-binding domains (DBDs) and additional carboxyl amino acids aligned using MUSCLE (Edgar 2004). The zinc fingers and basic domains are shown for representative sequences of (A) MED, (B) END-3, (C) END-1, and (D) a representative subset of all three factors. Consensus sequences are shown below each alignment. The phylogeny of Stevens *et al.* (2019) is shown to the left of the species names for reference. Under the consensus sequences, the amino acids that mediate site recognition by the *C. elegans* MED-1 DBD for (A) and cGATA1 for (B), (C) and (D) are shown (Lowry *et al.* 2009; Omichinski *et al.* 1993). Asterisks show corresponding amino acids that are invariant (black) or are generally conserved (gray).

In contrast with the divergent MEDs, the DBDs of the END-3 and END-1 orthologues are more alike and share greater similarity to those of canonical GATA factors. The ENDs, ELT-2 and cGATA have an invariant CXXC-X_17_-CXXC zinc finger structure with 17 amino acids between the 2^nd^ and 3^rd^ cysteines. Consensus sequences for END-3 and END-1, shown below the alignments in Figs. 9B and 9C, contain 23 invariant amino acids for END-3, and 31 for END-1, i.e. 2x and 3x more than the 11 invariant amino acids among the MED DBDs. A solution structure for END-1 or END-3 has not been reported, but as a surrogate I have shown, beneath both alignments, the 18 amino acids in the cGATA1 zinc finger known to mediate base contacts (Omichinski *et al.* 1993). END-3 is conserved at 7/18 of these positions with 4 amino acids being invariant, while END-1 has 10/18 positions conserved, of which 8 are invariant. Hence the END-1s are structurally more like cGATA1 than are the END-3s, plus the END-1 orthologues are also invariant at more positions, indicating that they are under the most evolutionary constraint.

An amino acid in the END-3 DBD is worth further comment. The proline between the 3^rd^ and 4^th^ cysteines of the zinc finger, in sequence CNPC, was substituted by a leucine in the EMS-induced *C. elegans* mutant *end-3(zu247)* (Maduro *et al.* 2005a). This mutant has a phenotype indistinguishable from the null mutant *end-3(ok1448)* which lacks most of the DBD (Owraghi *et al.* 2010). While this position is also a proline in 12/20 species, among the other END-3s it is serine (S) or alanine (A). Serine has a short polar side chain, while alanine is short and hydrophobic, however leucine is also hydrophobic but longer, suggesting that the longer side chain at this position compromises the structure of the zinc finger. This position is variable among the MED and END-1 orthologues, where it is a proline (P), alanine (A), serine (S), or glycine (G), indicating this position is under relaxed selection.

Another difference between the END-3s and END-1s is the amino end of the C4 zinc finger between the 1^st^ and 2^nd^ cysteines. GATA factors in general, including the MEDs, END-3, ELT-2 and cGATA1, have two amino acids in the pattern CXXC. Most of the END-3s are CSNC, while the END-1s have either CSNPNC (12 species), CSNPSC (6 species), CSNQNC (*C. afra*) or CNPNC (*C. becei*). It is not known what effect the extra one or two amino acids have on the structure of the zinc finger, however this variation in structure is found only in the END-1 orthologues.

Finally, as a set, the DBDs from the MEDs and ENDs of a subset of the Elegans supergroup species are shown with ELT-2 and cGATA1 in Fig. 9D, showing that all three factors share conserved amino acids with each other and with canonical GATA factors. Overall, 7/18 of the amino acids known to mediate DNA recognition in cGATA1 are broadly conserved (Omichinski *et al.* 1993).

### SERINE-RICH DOMAINS IN MEDs AND ENDs

The MED and END factors share an upstream region of variable size enriched in the polar amino acids serine, with or without threonine. These are shown diagrammatically in Fig. 8, as the amino-most conserved domain among the MEDs and ENDs, and in amino acid sequence alignment in Fig. 10A. Among the MEDs, the Poly-S/T region is variable in size, consists of both serines and threonines, and is the only other conserved feature upstream of the DNA-binding domain. Because of the size variability, the alignment in Fig. 10A represents only part of an overlapping region among MEDs of all 20 species. Among the ENDs, a similar Poly-S domain, consisting almost exclusively of homopolymeric clusters of serines, is found at the amino terminus starting at the 3^rd^ or 4^th^ amino acid (Fig. 10A). In one exception, the Poly-S domain is all but gone in *C. japonica* END-3. As noted earlier, the Poly-S region had been previously recognized in the *C. elegans* and *C. briggsae end* genes (Maduro *et al.* 2005a).

**Fig. 10.**
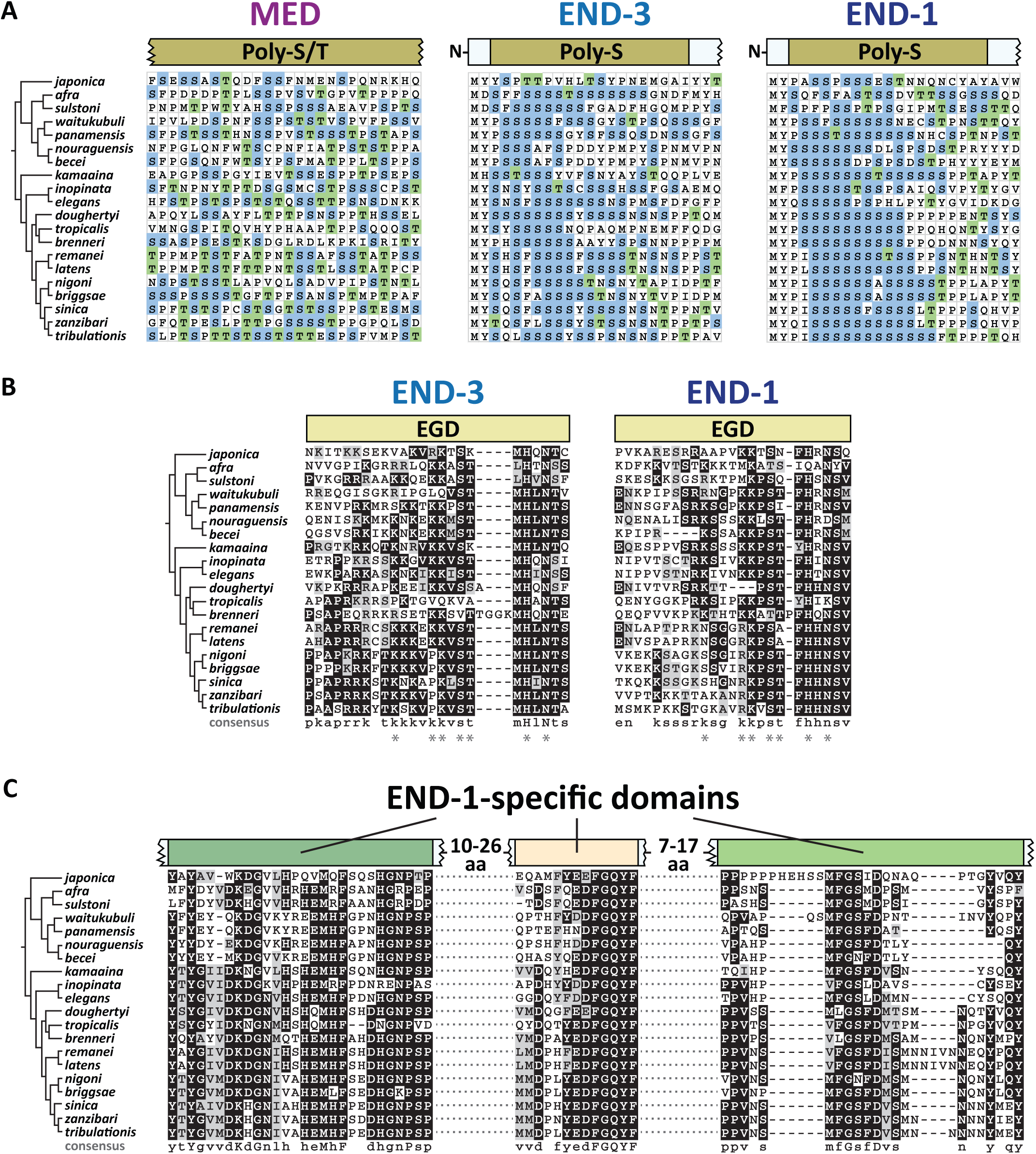
Other conserved domains of unknown significance among the MED and END proteins. (A) A portion of the alignment of Poly-S/T domains (MED factors) and the Poly-S domains (END-3 and END-1). Serines are highlighted in blue and threonines in green. (B) Extended Endodermal GATA Domains (EGDs) immediately upstream of the zinc fingers of END-3 and END-1. A consensus sequence is shown beneath each alignment, with amino acids similar between END-3 and END-1 shown with an asterisk (*). (C) Highly conserved regions among the END-1 factors showing highly conserved amino acids and a consensus sequence beneath the alignment.

An unexpected feature of the Poly-S region in the *end* genes bears further description. Although serine is coded by six codons – TCT, TCC, TCA, TCG, AGT and AGC – the serines among the Poly-S regions in the *end-3* and *end-1* orthologues are coded almost exclusively (99%, 554/557) by TCN codons (N=any base). Moreover, two of the four TCN codons, TCT and TCC, are used 50% and 22% of the time. Among *C. elegans* genes, TCN represents 75% of serine codons, and among these, TCT and TCC occur only 28% and 13% of the time, respectively (https://www.kazusa.or.jp/codon/). This preferential use of TCT and TCC codons for serine in the Poly-S regions, among the TCN codons, is statistically significant (p<10^−40^, χ^2^-test). The implications of this codon bias are discussed later.

### CONSERVATION OF THE END FAMILY GATA DOMAIN (EGD)

Previous work identified the END family GATA Domain, or EGD, immediately upstream of the *C. elegans* and *C. briggsae* END-1 and END-3 DBDs (Maduro *et al.* 2005a). This domain does not occur among the other *C. elegans* GATA factors, suggesting it is uniquely important for function of END-1 and END-3. Among the 20 species in the Elegans supergroup, the END-1 and END-3 orthologues across 20 species do contain a conserved region immediately upstream of the zinc finger. This is shown diagrammatically in Fig. 8, and by sequence alignment in Fig. 10B. Whereas the original report had the domain consisting of the 9 amino acids, an extended domain is apparent that consists of approximately 25 amino acids. 7 of these (shown by an asterisk in the figure) are highly conserved between the END-3 and END-1 factors, but there are conserved amino acids within each group of factors, plus the domain is more conserved among the END-3 orthologues. While the EGDs tend to be enriched in basic amino acids, their significance remains unknown.

### END-1 SPECIFIC DOMAINS

Among the END-3 orthologues, the region between the Poly-S and the EGD regions is variable in size and does not exhibit sequences with extensive conservation (Fig. 8). In contrast, the END-1 orthologues display three additional domains that are highly conserved across all 20 species (Figs. 8 and 10C). A consensus sequence shows high conservation with many invariant regions. These domains are apparently novel, as a BLAST search using this region of END-1 did not identify related proteins other than predicted orthologues of END-1 within *Caenorhabditis*. With the identification of these extended sequence similarities, the END-1 orthologues across the 20 species are highly conserved throughout their lengths, while the END-3 and MED orthologues are conserved only in parts.

## Discussion

In this work I have identified and compared the gene and protein structures of the MED, END-3 and END-1 GATA transcription factors among 20 *Caenorhabditis* species of the Elegans supergroup. Predictions were made by manual curation, informed by known features of the network from *C. elegans* and informed by comparison of gene and protein structures together. The results confirm coevolution of *cis*-regulatory sites, gene structures and protein sequence over tens of millions of years of evolution. Many of the conserved features, including the DNA-binding domains, and binding sites for SKN-1, MED, and an Sp1-like factor, are consistent with known properties of the *med* and *end* genes in *C. elegans* (Broitman-Maduro *et al.* 2005; Maduro *et al.* 2015; Maduro *et al.* 2001; Sullivan-Brown *et al.* 2016). Prior work has also shown that orthologous *med*s and/or *end*s from a few of these species can function as transgenes in *C. elegans* (Coroian *et al.* 2005; Maduro *et al.* 2005a). Hence, I hypothesize that the *med*, *end-3* and *end-1* genes function in a core endoderm specification network across the Elegans supergroup that originated in a common ancestor.

### HIGH RATES OF MED GENE DUPLICATION

The *med*, *end-3* and *end-1* genes showed distinct patterns of gene duplication among species. Occurrence of duplicate *med* genes is disproportionately high, with an average of 5.6 *med* genes per species, compared with 2.0 *end-3* genes and a single *end-1* per species, except for *C. brenneri* which may have two *end-1* genes (Fig. 2). In most cases, sequence similarity was consistent with most *med* duplicates having arisen post-speciation, with the only exceptions resulting from likely inheritance of two *med* genes in a recent common ancestor (Fig. 6).

The apparent recent amplification of the *med*s suggests that there is ongoing selective pressure for increased *med* expression. The occurrence of MED binding sites in the *end* genes (particularly *end-3*) argues for positive selection for the presence of these sites, and hence the MED factors that can bind them. Selection for increased *med* expression is supported by work showing that *C. elegans* has an unusually high rate of segmental duplications compared with other species, with a higher gene dose generally leading to increased mRNA production (Konrad *et al.* 2018). In *C. elegans*, a single chromosomal *med* gene is sufficient for completely normal development (Maduro *et al.* 2007). However, *C. elegans* has only two *med* genes. Perhaps in some of the other species, the MED factors have become degenerate in their ability to activate target genes, or to be activated. Protein degeneracy would be consistent with the lower degree of protein sequence conservation among the MED DNA-binding domains in *C. brenneri*, which has experienced an extreme amplification of *med* genes (Fig. 9). However, that does not explain amplification of *med* genes in *C. doughertyi*, whose MED DNA-binding domains are more similar as a group, unless they are collectively degenerate in some way (Fig. 9). Regardless of the mechanism driving MED amplification, there is support for reduced fitness if MED-dependent input into endoderm specification is compromised. Recent work has found that loss of MED binding sites in the *end* genes in *C. elegans* results in aberrant intestinal lineage development, metabolic defects, and reduced viability (Choi *et al.* 2017; Maduro *et al.* 2015). Another possibility, not mutually exclusive, is that degeneracy of MED function leads to embryonic lethality due to a failure to specify the MS blastomere (Maduro *et al.* 2001). Hence, whatever mechanism driving is increased *med* dosage may not be due to the role of the MEDs in gut specification.

### LINKAGE OF END ORTHOLOGUES

In most species, *end-1* was found within ~35 kbp of *end-3* (Fig. 3A). One possibility for maintenance of this synteny is that the two genes may be coregulated. Three lines of evidence argue against this possibility, at least for *C. elegans*. First, there is at least one unrelated gene between the *end*s, the neural gene *ric-7* (Hao *et al.* 2012). Second, the *end-1,3* genes are not precisely co-expressed as accumulation of *end-3* mRNA precedes that of *end-1* (Baugh *et al.* 2003; Maduro *et al.* 2007; Raj *et al.* 2010). Third, unlinked single-copy transgenes of wild-type *end-1* and *end-3* are able to completely replace function of the endogenous genes when introduced into an *end-1,3(-)* strain, suggesting that linkage is not a prerequisite for their expression (Maduro *et al.* 2015). It may be, therefore, that synteny of *end-1* and *end-3* merely reflects their origin as a tandem duplication of an ancestral *end* gene.

### IDENTIFICATION OF KNOWN AND PREVIOUSLY UNRECOGNIZED *cis*-REGULATORY SITES

The MEME search recovered binding sites for regulators previously known to activate the *med* and *end* genes in *C. elegans* (Fig. 4B). In the case of the *med* orthologues, this was binding sites for SKN-1, while for the *end* genes, it was binding sites for both SKN-1 and MED-1. The conservation of these sites supports the hypothesis that these genes have maintained the same regulatory hierarchy as in *C. elegans*, with SKN-1 activating the *med* genes, and both SKN-1 and the MED proteins activating the *end* genes. The MED sites in the Elegans supergroup *end* genes are found in all *end-3* orthologues but only 9/20 *end-1* orthologues, following the same pattern as in *C. elegans*: *end-3* has four MED sites and these are collectively essential for *end-3* activation, although even a single MED site in a single-copy *end-3* transgene is sufficient for activation (Maduro *et al.* 2015). In contrast, *end-1* has only two MED sites, and these are less important for *end-1* expression due to parallel input by TCF/POP-1 and PAL-1 (Maduro *et al.* 2015; Maduro *et al.* 2005b). The likely sites for SKN-1 in *end-1* and *end-3* were not previously known because they do not contain the same pattern of SKN-1 site core sequences as present in the *med* promoters. An intriguing hypothesis is that the SKN-1 sites in the *end* genes may be of lower affinity than those in the *med* genes. Because expression of the *end* genes is delayed by at least one cell cycle compared with *med-1,2*, lower-affinity SKN-1 sites could potentially allow for delayed activation. A similar affinity difference has been hypothesized for early- and late-acting binding sites of the pharynx regulator PHA-4 (Gaudet *et al.* 2004). As the SKN-1 sites in the *end* genes were not found in all species, it is possible that the input from SKN-1 is lost in some species. Finally, an additional suspected regulatory input was from an Sp1-like factor, likely to be SPTF-3 (Sullivan-Brown *et al.* 2016). Most of the *med*, *end-3* and *end-1* orthologues have a consensus Sp1 binding site (Fig. 4B). Together, the recovery of these sites from an *ab initio* search of their putative promoters lends strong support to the hypothesis of conservation of this gene network across the Elegans supergroup.

MEME-identified sites of lower significance, and not as broadly conserved, were either unknown or reflected putative core promoter elements. These include one with core sequence TCTKCAC, a polypyrimidine motif, putative PolyA/T cluster, a TATA-binding protein (TBP) site, and an SL1 motif. The latter two were previously found in many promoters in five Elegans supergroup species (Grishkevich *et al.* 2011). The putative PolyA/T cluster is associated with germline expression (Frokjaer-Jensen *et al.* 2016). The other two motifs were of unknown significance. The TCTKCAC motif is found in *C. elegans med* genes, hence it is possible to test its significance directly. The site was found three times close to the previously identified SKN-1 sites, suggesting it may play an accessory role to SKN-1 activation, perhaps by SKN-1 itself.

What was particularly conspicuous was that sites for minor regulatory inputs known in *C. elegans* were not found to be widely conserved, either by a direct search or through MEME. This includes sites for TCF/POP-1 and the Caudal orthologue PAL-1, both of which are genetically known to contribute to *end-1* expression, and for which binding sites are known or suspected based on prior work (Bhambhani *et al.* 2014; Maduro *et al.* 2005b; Robertson *et al.* 2011; Shetty *et al.* 2005). *C. elegans* END-3 is also a suspected contributor to activation of *end-1* (Maduro *et al.* 2007). The failure to recover sites for these regulators suggests that either these inputs exist in the other species and are not recognizable, or more likely, that different species have qualitatively different minor regulatory inputs. The apparent difference in regulatory input of SKN-1 and POP-1 in *C. briggsae*, revealed through cryptically different reduction-of-function phenotypes between *C. briggsae* and *C. elegans*, suggests that reinforcing regulatory inputs may evolve rapidly (Lin *et al.* 2009). Even within *C. elegans*, widespread cryptic variation in input from SKN-1 and the Wnt pathway (which acts through POP-1) was observed among *C. elegans* wild isolates (Torres CLEUREN *et al.* 2019). An emerging model seems to be that the core SKN-1 → MED → END-1,3 regulatory cascade is conserved, while additional regulatory inputs that reinforce this cascade evolve rapidly and would thus be expected to be species-specific. Putative *cis*-regulatory sites that mediate these supporting inputs might therefore occur in only a subset of species in the Elegans supergroup and would be missed in the analysis done here.

### END-3 AND END-1: THE SAME BUT DIFFERENT

In *C. elegans*, *end-1* and *end-3* clearly have overlapping function. Complete loss of both genes has a fully penetrant failure to specify endoderm, while null alleles either for gene alone have either no effect (*end-1*) or a weak effect (*end-3*) on gut specification (Maduro *et al.* 2005a; Owraghi *et al.* 2010). A similar result was obtained using RNAi in *C. briggsae* (Maduro *et al.* 2005a). As well, overexpression of either *end* gene in *C. elegans* is sufficient to induce endoderm differentiation in non-endodermal lineages (Maduro *et al.* 2005a; Zhu *et al.* 1998). Within their DNA-binding domains, the END-3 and END-1 orthologues are clearly more similar to each other than they are to the MEDs (Figs. 5, 9).

Despite these similarities, END-3 and END-1 differ in ways that suggest they have at least some unique functions. First, the END-1 DBDs are more highly conserved as a group, while those of END-3 are under slightly more relaxed selection. This is apparent in the way that the DBDs appear in a phylogenetic tree (Fig. 7) and in the degree of invariant amino acids in an alignment (Figs. 9B, 9C). Within their DBDs, the END-1s have twice as many similar amino acids in common with vertebrate cGATA1 than the END-3s have in common with cGATA1, notably in acid positions known to mediate sequence recognition (Figs. 9B, 9C).

Additional evidence is consistent with both shared and divergent activity of END-3 and END-1 in *C. elegans*. Recent work inferred the binding sites for *C. elegans* END-1 and END-3 as RSHGATAASR and RKWGATAAGR, respectively, which are very similar though not identical (Lambert *et al.* 2019; Weirauch *et al.* 2014). Other work has shown that recombinant DNA-binding domains of *C. elegans* END-1 and END-3 can bind canonical GATA sites in the promoter of *C. elegans elt-2*, although END-1 has a higher affinity for such sites (Du *et al.* 2016; Wiesenfahrt *et al.* 2015). From this work, Endoderm GATA Domains (EGDs) immediately upstream of the DBDs show conserved amino acids between END-3s and END-1s but many more that are unique to either EGD (Fig. 10B). Although the function of the EGDs remains unknown, their conservation and proximity to the DBDs suggest an accessory role in protein-DNA interactions that is unique to the ENDs among the *Caenorhabditis* GATA factors.

### THE POLY-S REGION OF END-3 AND END-1: PROTEIN DOMAIN OR POLYPYRIMIDINE TRACT?

END-3 and END-1 share an amino-terminal segment, far from the DNA-binding domain, that is enriched for homopolymers of serine (Fig. 10A). Such a domain is not found in the other *C. elegans* GATA factors, nor is enrichment for serine found in vertebrate GATA factors (Kaneko *et al.* 2012; Yang *et al.* 1994). This suggests that the Poly-S domain plays some other function besides DNA binding and transactivation. The selection for TCT and TCC codons suggests that the Poly-S regions have been maintained for a reason other than a selection for what they contribute to the END-1 and END-3 proteins. Beyond transcriptional activation of the *end-1* and *end-3* genes, post-transcriptional regulatory mechanisms could potentially fine-tune END-1,3 protein levels. At the level of mRNA, the preference for these codons, as opposed to UCG and UCA, results in maintenance of a polypyrimidine tract in the mRNA. Support for a possible role of such a tract in the endoderm GRN is that in some species (e.g. *C. latens* and *C. remanei*), the *med* orthologues also have an apparent enrichment of T and C bases in the first part of their coding regions. In other systems, polypyrimidine tract binding proteins (PTBs) have various roles in RNA metabolism, including regulation of splicing and mRNA stability, though in these cases the tracts occur outside of coding regions (Sawicka *et al.* 2008). There is a *C. elegans* PTB gene, *ptb-1*, but its function has not been described. At the level of translation, repeats of the same UCY serine codon could cause starvation for limiting amounts of a particular seryl-tRNA^Ser^, leading to ribosome pausing (Darnell *et al.* 2018). However, it is not clear why there would be selection to delay translation of *end* mRNA, particularly as given the rapid early cell divisions of the *C. elegans* embryo, it makes more sense to express the gene products as rapidly as possible. A more benign reason for the maintenance of the serine codon repeats is that they are an artifact of a trinucleotide repeat expansion process (Koren and Trifonov 2011). Indeed, in that study, amino acid repeats in vertebrate proteins were most likely to be found in the first exon, i.e. at the amino end, consistent with their location in the *end-3* and *end-1* genes. Hence, the role of the Poly-S domain, if any, remains open for speculation until structure-function studies are performed.

### END-1 ORTHOLOGUES ARE CONSERVED THROUGHOUT THEIR LENGTHS

An additional unexpected finding emerged from the alignment of END-1 orthologues that distinguishes them among the MED/END proteins. Between the Poly-S and EGD domains, the END-3 orthologues as a group were diverse in size and sequence, whereas the END-1 orthologues were more similar in size and showed several regions of high conservation (Fig. 10C). These END-1-specific domains could be grouped into three regions containing blocks of invariant amino acids. The most striking of these is the center domain which contains an invariant sequence of FGQYF across all species END-1s. None of these highly conserved domains are found in other proteins, apart from predicted END-1 orthologues. The high conservation is further supported by the conservation of introns. The END-1s have four introns with only one of these absent in *C. brenneri* (Fig. 4A). In contrast, the END-3s were more likely to experience intron gains and losses over the same evolutionary time period, with most of these occurring in the variable region between the amino-terminal Poly-S and EGD domains (Fig. 8). A cursory examination of the amino acids in the END-1-specific domains suggests that these are on the outside of the protein, perhaps mediating protein-DNA or protein-protein interactions that do not occur with END-3 (data not shown).

Taken together, these data show that across the Elegans supergroup, the END-1s are highly conserved proteins with greater similarity to vertebrate GATA factors than the more diverse END-3s paralogues. This predicts that END-1 has unique features in transcriptional activation, and that the target genes activated by each of these factors are likely to include both and distinct targets.

### MED ORTHOLOGUES: A DIVERGENT AND DIVERSE SUBCLASS OF GATA FACTORS

The MED orthologues among the 20 species were found to be divergent from the END-3/END-1 factors, and to comprise a more diverse group of proteins themselves, even within the DNA-binding domain (Figs. 5, 9). The divergence of the DBD from that of the ENDs, ELT-2 and cGATA is expected, because the *C. elegans* MEDs were recognized to be divergent GATA factors that recognize a different binding site with an AGTATAC core (Broitman-Maduro *et al.* 2005; Lowry *et al.* 2015). Despite the high divergence of the MED factors as a group, indicating relaxed selection, there is nonetheless maintenance of their binding site sequence over evolutionary time. This is supported by the conservation, across all 20 species, of most of the amino acids that were found to mediate protein-DNA recognition in *C. elegans* MED-1 (Fig. 9A), and more importantly, by the MEME identification of AGTATAC binding sites among all *end-3* orthologous genes and 9/20 *end-1* genes (Fig. 4). Furthermore, transgenes of most of the *C. briggsae* and *C. remanei med*s were individually able to complement *C. elegans med-1,2* double mutants in both gut and mesoderm specification despite limited conservation (Coroian *et al.* 2005). Selection is likely not acting solely on the MEDs for *end* gene activation, as there are other direct MED targets in *C. elegans* whose orthologues in the Elegans supergroup were not investigated here, including in the early MS lineage (Broitman-Maduro *et al.* 2006; Broitman-Maduro *et al.* 2005). The lower conservation suggests that the MED DBDs may simply be more accommodating of amino acid substitutions than are the DBDs of END-3 or END-1.

Outside of the DNA-binding domain, the MEDs as a group lack the type of conserved regions seen in the ENDs. The only other feature found is a variable enrichment for serine and threonine of unknown significance. This region does not resemble the homopolymeric enrichment for serine that is at the amino end of the ENDs (Fig. 10A). Rather, it is a higher prevalence for S/T that lacks a recognizable context. A serine-threonine rich motif was found to be important for nuclear localization of the mineralocorticoid receptor in vertebrates, suggesting that this region of the MED orthologs may play a similar role (Walther *et al.* 2005). Until structure-function analyses are done, the significance of the serine/threonine enrichment will remain unknown.

### THE MED/END CASCADE IS A DERIVED CHARACTER

The existence of a gut-like precursor is a conserved lineage feature found in more distantly related nematode species (Houthoofd *et al.* 2003; Schierenberg 2006; Schulze and Schierenberg 2011). It must therefore be that species outside the Elegans supergroup specify the gut precursor without MED/END factors. The most upstream factor SKN-1, and the downstream gut identity factor ELT-2, are also more widely conserved than just the Elegans supergroup (Couthier *et al.* 2004; Schiffer *et al.* 2014). Assuming that SKN-1 still specifies MS and E, the simplest hypothesis is that specification of gut outside of the Elegans supergroup occurs by direct activation of an *elt-2*-like gene directly by SKN-1. An attempt to demonstrate bypass of the *end-1* and *end-3* genes was successful using an *elt-2* transgene under regulatory control of the *end-1* promoter in a *C. elegans* strain lacking *end-1* and *end-3* (Wiesenfahrt *et al.* 2015). However, this transgene worked best in a high copy-number array, and not in single-copy. Furthermore, expression of this transgene is likely to be at least partially dependent upon regulatory input by MED-1,2, based on studies with an *end-1* promoter lacking MED binding sites (Maduro *et al.* 2015). As an alternative to direct SKN-1 → ELT-2 regulation, there could be one or more non-GATA regulators between them, analogous to the MED/END cascade. Regardless of how gut specification occurs outside of the Elegans supergroup, some set of evolutionary events must have set in motion a breakdown of the ancestral specification mechanism, favoring the evolution and fixation of the SKN-1/MED/END cascade as the dominant mode of E specification.

### EVOLUTIONARY ORIGIN OF THE SKN-1 → MED → END-1,3 CASCADE

The co-occurrence of the MED and END factors suggests that these genes evolved within a short time at the base of the Elegans supergroup (Fig. 11A). At the start of this work there was an expectation that there might have been one or more "transitional" species with only the *end-3* and *end-1* factors, or only one *end-*like factor, for example. Since no such species were found, it may be that a transitional species has not yet been sequenced, or that the orthologues are highly diverged. The reduced number of recognizable GATA factors in species outside of the Elegans supergroup argues against this possibility, however.

**Fig. 11.**
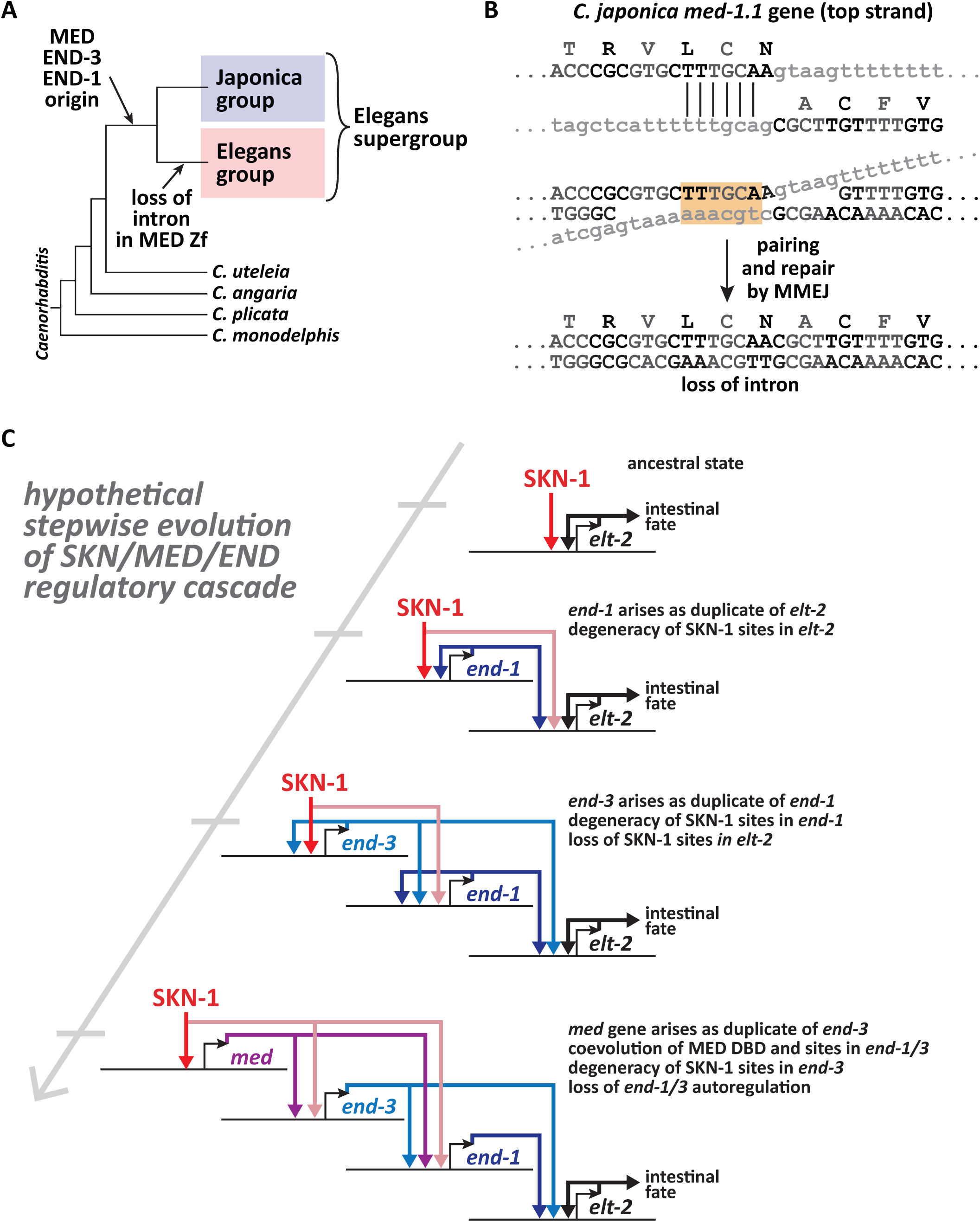
Origin of the MED, END-3 and END-1 factors. (A) Origin of all three factors at the base of the Elegans supergroup, followed by loss of a conserved intron in an ancestral *med* gene at the base of the Elegans group. (B) Hypothetical microhomology-mediated end joining (MMEJ) event that could delete the conserved zinc finger intron at the base of the Elegans group, using a 6-bp identity in-frame microhomology in an extant *C. japonica med* gene. At top, the microhomology is shown for the top strand. In the bottom part, complementary strands are shown pairing across the microhomology, which if resolved could result in an in-frame deletion of the intron, after (van Schendel and Tijsterman 2013). This would also require maintenance of the AAC codon for asparagine immediately to the right of the homology. (C) Speculative model for generation of the SKN-1/MED/END regulatory cascade through intercalation by serial duplications of an ancestral autoregulating *elt-2* gene. A bent arrow indicates the transcription start site, with the regulatory activity of the protein product of the gene shown as a colored line from the bent arrow. The promoter is to the left of the bent arrow. The positions in the promoters are only meant to qualitatively convey positive regulation and not indicate number or position of binding sites.

The data strongly suggest that the *med* and *end* genes might have been derived from the same ancestral gene. This hypothesis is supported by the existence of an intron in the zinc finger domain of all *med* and *end* genes, except for the Elegans group *med* genes where loss of this intron occurred. In the genus, intron loss is common, and occurs more frequently than intron gain (Roy and Penny 2006). One mechanism by which this particular intron could have been lost is through germline gene conversion from a reverse-transcribed (spliced) mRNA (Roy and Gilbert 2005). An alternative mechanism could be through microhomology-mediated end joining, or MMEJ, of a double-stranded break in the gene (Mcvey and Lee 2008; van Schendel and Tijsterman 2013). Indeed, in one of the *C. japonica med* genes, a short stretch of six base pairs upstream of this intron recurs close to the 3’ splice site of the intron itself, such that a repair of a double-stranded chromosome break by MMEJ would result in an in-frame removal of the intron (Fig. 11B). This would also require that the asparagine codon (AAC) is somehow maintained, which may be possible given the observed types of MMEJ repair of double-stranded breaks induced by Cas9 cleavage, e.g. (Taheri-Ghahfarokhi *et al.* 2018). Regardless of the mechanism, loss of this intron likely occurred only once in the last common ancestor to the Elegans group. I note in passing that the converse property, lack of intron gain in the Elegans group *med* genes, may be accounted for by selection for rapid gene expression through avoidance of mRNA splicing; most early zygotic *Drosophila* genes are in fact intronless (Guilgur *et al.* 2014). However, a small number of the *med* gene predictions in the Elegans supergroup do have introns (Supplemental File S1).

The structural conservation among the 20 Elegans supergroup MEDs and ENDs lead me to propose a model by which the MED/END cascade arose through duplication and modification of existing genes, from *elt-2* upwards, as shown in Fig. 11C. The similarity of the END-3 and END-1 orthologs and their tendency to be <50 kbp apart in a species suggests that they originated from a common progenitor together, or that one was a duplicate of the other. Considering the stronger resemblance of the DNA-binding domain of END-1 with that of ELT-2 and vertebrate cGATA1, a reasonable hypothesis is that *end-1* originated first, as a duplicate of an ancestral *elt-2* gene that was both activated by SKN-1 and maintained its own expression through positive autoregulation. Positive autoregulation of ELT-2 is known and has even been visualized *in vivo* (Fukushige *et al.* 1999). Duplication of *elt-2* has likely occurred to generate the extant paralogous (and likely inactive) *C. elegans elt-4* gene, and more significantly, *C. elegans elt-7*, a paralogue of *elt-2* that shares overlapping function, expression and autoregulation with *elt-2* (Fukushige *et al.* 2003; Sommermann *et al.* 2010). Although not necessary at this step, if the SKN-1 sites in the *elt-2* promoter became degenerate, the *end-1* prototype would be stable. A paralogous *end-3* prototype gene might then have originated as a simple linked duplication of *end-1*. Lending support for *elt-2* as a progenitor for the *end* genes is the presence of the conserved zinc finger intron found in all *end-1/3* orthologues and in *C. elegans elt-2*/*7*. The two *end* genes could be stabilized by the complete loss of SKN-1 sites in the *elt-2* promoter, degeneracy of SKN-1 sites in the *end-1* promoter, and coevolution of END-3 with binding sites in the *end-1* promoter. In this state, *end-1* acts to amplify input into *elt-2* from *end-3*.

A challenge is in accounting for the origin of a *med*-like progenitor, given the evidence that they form a structurally divergent set of regulators. In this work it was found that while the Elegans group species have intronless *med* genes, obscuring their origin, the putative Japonica group *med*s share a common intron in the zinc finger coding region that is in the same location as the aforementioned intron in all extant *end-3* and *end-1* genes. This leads to the hypothesis that a prototype *med* gene arose as a duplicate of one of these genes, the most logical of which may be *end-3*. Co-evolution of the MED DNA-binding domain with cognate sites in *end-1* and *end-3* would reduce autoregulation of the *end* genes and fix the MED factor within the network, though END-3 could retain the ability to contribute to *end-1* activation. Degeneration of the SKN-1 sites in *end-3* would strengthen the feed-forward cascade. Further refinement of the network would strengthen regulatory input of the *med*s by SKN-1, activation of *end-3* by the MEDs, and other regulatory inputs into *end-1*. Further selection on the END-1 coding region might have been enforced by protein-protein interactions with other factors that contribute to gut specification.

Although this model is highly speculative, there is supporting evidence from evolution of the *Bicoid* (*Bcd*) gene in an ancestor to cyclorrhaphan flies, a group that includes *Drosophila* (Driever and Nusslein-Volhard 1989; Stauber *et al.* 1999). *Bcd* specifies anterior fates in early cyclorrhaphan embryos, while outside of this group *bcd* is not found, and other factors play an analogous role (Lynch *et al.* 2006; Mcgregor 2005). *Bcd* arose as a duplicate of the Hox gene *Zen*, and likely acquired derived DNA-binding characteristics primarily through two missense mutations in the DNA-binding domain (Liu *et al.* 2018; Mcgregor 2005). From studies in the flour beetle *Tribolium*, which lacks *bcd*, it is hypothesized that *Bcd* took over functions of some of its downstream gap gene targets, which it then became an activator of (Mcgregor 2005). *Bcd* is proposed to have originated ~140 Mya at the base of the Cyclorrhapha, a longer time period than the estimated tens of millions of years since the common ancestor to the Elegans supergroup (Coghlan and Wolfe 2002; Cutter 2008; Wiegmann *et al.* 2011). Recruitment of *Bcd* into A/P specification in *Drosophila* likely required more steps than the MED/END cascade, because from the proposed model, the cascade originated through duplication and modification of a factors already in an ancestral version of the network. Hence, it is plausible that emergence of the MED/END network could have occurred at the base of the Elegans supergroup. Furthermore, in analogy to *Bcd*, the initial evolution of the MED DBD that resulted in a change in its binding site to a non-GATA target site might have been driven by a small number (or even just one) key amino acid change. With the sequences of *med* genes from 20 species, such structure-function correlations can now be examined.

Studies on the evolution of *Bcd* suggest a possible explanation as to why a more layered gene cascade might have evolved for embryonic gut specification within the Elegans supergroup. The emergence of *Bcd* may have conferred a more rapid specification of segment identity, allowing developmental time to become faster without sacrificing robustness (Mcgregor 2005). By extension to the Elegans supergroup, it is possible that the SKN-1 → MED → END-1,3 gene regulatory cascade coincided with an increase in developmental speed in *Caenorhabditis*, perhaps as part of the transition to very early and rapid cell fate specification (Laugsch and Schierenberg 2004; Schierenberg 2001). Elucidation of gut specification mechanisms in *Caenorhabditis* species outside of the Elegans supergroup, compared with their developmental speed, could provide evidence for this hypothesis, or alternatively identify non-GATA factors that play the same role as the MED/END cascade.

In the meanwhile, the identification of MED, END-3 and END-1 orthologues in 20 species sets the stage for studies to test hypotheses about evolution of gene regulatory networks, structure-function correlations in the evolution of novel DNA-binding domains, and features of developmental system drift. As the study of gene regulatory networks becomes more computational, the set of MED and END orthologues identified here will provide a basis for future studies integrating gene network architecture with transcriptomics data, for example (Nomoto *et al.* 2019; Omranian and Nikoloski 2017).

## ACKNOWLEDGMENTS

I am indebted to Mark Blaxter, Lewis Stephens and colleagues at the *Caenorhabditis* Genomes Project in Edinburgh for prepublication access to the genome sequences of *Caenorhabditis* species and for advice during this work. I also thank Eric Haag (University of Maryland, College Park) for helpful advice in interpretation of search results. Earlier versions of this work were completed under my NSF grant IOS#1258054. I also thank Christian Turner, former UCR undergraduate supported by NIH Award T34GM062756 from the National Institute of General Medical Sciences (MARCU-STAR) program to UC Riverside, for having performed preliminary searches of available *Caenorhabditis* sequences in 2014.

---

**Supplemental File S1.** This Microsoft Excel (.xlsx) file contains all gene predictions, coding regions, and coordinates of protein domains used to generate Fig. 8.

**Supplemental Files S2-S12.** FASTA files containing protein and promoter sequences.

**Supplemental Files S13 and S14.** MEME output HTML files.

**Supplemental Tables S1, S2 and S3.** These tables contain search results for known *cis*-regulatory sites.

